# Strigolactone regulates shoot development through a core signalling pathway

**DOI:** 10.1101/070763

**Authors:** Tom Bennett, Yueyang Liang, Madeleine Seale, Sally Ward, Dörte Müller, Ottoline Leyser

**Affiliations:** Sainsbury Laboratory, University of Cambridge, Bateman Street, Cambridge, CB2 1LR, UK; Current Address: Department of Biology, University of Leeds, Leeds, LS2 9JT, UK; Current Address: Institute of Molecular Plant Sciences, University of Edinburgh, Edinburgh, EH9 3BF, UK; Department of Biology, University of York, York, YO10 5DD

## Abstract

Strigolactones are a recently identified class of hormone that regulate multiple aspects of plant development. The DWARF14 (D14) α/β fold protein has been identified as a strigolactone receptor, which can act through the SCF^MAX2^ ubiquitin ligase, but the universality of this mechanism is not clear. Multiple proteins have been suggested as targets for strigolactone signalling, including both direct proteolytic targets of SCF^MAX2^, and downstream targets. However, the relevance and importance of these proteins to strigolactone signalling in many cases has not been fully established. Here we assess the contribution of these targets to strigolactone signalling in adult shoot developmental responses. We find that all examined strigolactone responses are regulated by SCF^MAX2^ and D14, and not by other D14-like proteins. We further show that all examined strigolactone responses likely depend on degradation of SMXL proteins in the SMXL6 clade, and not on other proposed proteolytic targets. Taken together, our results suggest that in the adult shoot, the dominant mode of strigolactone signalling is D14-initiated, MAX2-mediated degradation of SMXL6-related proteins. We confirm that the BRANCHED1 transcription factor and the PIN-FORMED1 auxin efflux carrier are plausible downstream targets of this pathway in the regulation of shoot branching, and show that BRC1 likely acts in parallel to PIN1.

**AUTHOR SUMMARY:** Strigolactones are a recently discovered family plant hormones with diverse roles in development, most strikingly in the regulation shoot branching. Our understanding of the mechanism(s) by which plants perceive and respond to strigolactones is growing rapidly. It is likely that the strigolactone signaling pathway has evolved by duplication and diversification of specific components of a pre-existing pathway, involved in perception and response to an as yet unknown hormone. Several of these components have been identified and several new candidate components have been implicated in the pathway. We have adopted a genetic approach to assess systematically the contributions of all these players to strigolactone signaling in the shoot. We exclude some of the candidate proteins from involvement in strigolactone-mediated shoot branching control and define a core pathway for strigolactone action in the shoot. We provide evidence that downstream of this core, the strigolactone signaling pathway branches, with different effectors mediating different shoot responses.

## INTRODUCTION

Plant development is a continuous process that is modulated by multiple environmental stimuli. Many of these stimuli are perceived locally, but require global and/or systemically co-ordinated responses. A small number of low molecular weight signalling molecules, including auxin and cytokinins, have been implicated in this intra-plant communication. Of these signals, the most recently identified are the strigolactones (SLs), a group of carotenoid-derived terpenoid lactones. Strigolactones (SLs) were first identified as a component of root exudates that cause seed germination in parasitic witchweeds (*Striga* spp.) (reviewed in Xie et al, 2010). Subsequently, root exudation of SL was shown to be required for the establishment of symbioses with arbuscular-mycorrhizal (AM) fungi, a process which has been hijacked by parasitic plants (Xie et al, 2010). In parallel, genetic and physiological studies in several species suggested the existence of a carotenoid-derived long-distance endogenous signal, which was subsequently shown to be SL (Gomez-Roldan et al, 2008; Umehara et al, 2008). Mutation in SL signalling and synthesis components confers a range of developmental phenotypes such as changes in shoot and root branching and elongation. Thus in higher plants, SLs function both as rhizosphere inter-organism signals and systemic intra-organism signals. These two distinct facets of SL function can be conceptualized as an integrated nutrient deficiency response, which is particularly related to nitrate and phosphate availability (Kohlen et al, 2011; Foo et al, 2013; Sun et al, 2014; de Jong et al, 2014). SL, primarily produced in the root, coordinates plant responses to nutrient deficiency by attracting AM fungi (which provide nutrients in return for fixed carbon), and remodelling the root and shoot systems, adapting growth to available resources.

SLs are synthesised by the action of at least 4 enzyme classes: the DWARF27-class carotenoid isomerases, the carotenoid cleavage dioxygenases CCD7 and CCD8 and the MAX1 (MORE AXILLARY GROWTH1)-class cytochrome P450s (reviewed in Waldie et al, 2014). The combined action of DWARF27 (D27), CCD7 and CCD8 produces carlactone, a MAX1 substrate which appears to be a precursor for a range of biologically active SLs identified in plants (Alder et al, 2012; Seto et al, 2014; Abe et al, 2014).. This core pathway is responsible for most SL synthesis, but plants lacking any one of these enzymes still produce some SLs, indicating that our knowledge of SL synthesis is incomplete (Waldie et al, 2014). Recent work suggests that there are likely to be multiple additional enzymes responsible for the further processing of carlactone into various active SLs (Brewer et al, 2016). Much recent progress has been made in understanding SL signalling (reviewed in Bennett & Leyser, 2014; Waldie et al, 2014). Genetic screen have identified two major classes of protein required for SL perception, namely the DWARF14-class of α/β-fold hydrolase proteins (Arite et al, 2009; Hamiaux et al, 2012) and the MAX2 class of F-box proteins (Stirnberg et al, 2002; Stirnberg et al, 2007). There is now very good evidence that D14 proteins act as strigolactone receptors, by cleaving of SLs and covalently retaining one of the hydrolysis products. This causes a conformational change in D14 that allows its interaction with MAX2 (de Saint Germain et al, 2016; Yao et al, 2016). MAX2 forms part of a Skip1-Cullin-F-box (SCF) E3 ubiquitin ligase complex (Stirnberg et al, 2007). Such complexes typically trigger the degradation of target proteins via the 26S proteasome, and have previously been demonstrated to be involved in many plant signalling pathways (Vierstra, 2009).

Intriguingly, MAX2 has also been implicated in responses to smoke-derived signalling molecules known as karrikins, which promote germination in fire-following species and share structural properties with SLs (Nelson et al, 2011). Karrikins also promote germination in non-fire following species such as Arabidopsis, leading to suggestions that exogenous karrikins piggyback on the signalling pathway of an as-yet-unidentified endogenous karrikin-like signalling molecule (Flematti et al, 2013), hereafter referred to as KL (Soundappan et al, 2015). The similarities between SL and KL signalling run deeper, since the receptor for KL, KARRIKIN INSENSITIVE2 (KAI2), is a close relative of D14 (Waters et al, 2012a). There is also a third member of the KAI2/D14 family, D14-LIKE2 (DLK2), which is highly conserved in flowering plants, but has no identified function (Waters et al, 2012a). Phylogenetic analysis suggests that D14 and DLK2 are recent innovations, arising in the vascular plant lineage, whereas KAI2 homologues are present throughout land plants and their algal relatives (Delaux et al, 2012; Waters et al, 2015). SLs are also present throughout the land plants and in some algae (Delaux et al, 2012). Moss mutants deficient in SL synthesis have colony extension defects, and the rhizoids of charophyte algae have been shown to respond to treatment with SL analogues, concordant with the idea that SLs are nutrient deficiency signals (Delaux et al, 2012; Proust et al, 2012). Though present in moss genomes, MAX2 does not appear to be involved in SL responses in *Physcomitrella patens* (de Saint Germain et al, 2013a), and these plants lack apparent D14 orthologues (Waters et al, 2015), suggesting that there may be alternative, more ancient SL signalling pathways present in basal land plants (Challis et al, 2013; Bennett & Leyser, 2014). For instance, some of the KAI2-like proteins present in the Physcomitrella genome appear to have binding pockets that could accommodate SLs, and might therefore be involved in SL perception (Lopez-Obando et al, 2016).

Since both SL and KL act through MAX2-dependent signalling, a goal in elucidating their mechanism of action is to identify the proteins marked for degradation by SCF^MAX2^, and determine whether there are common or separate targets of SL and KL signalling. Mutants in *SUPPRESSOR OF MAX2 1* (*SMAX1*), encoding a HEAT SHOCK PROTEIN101-like protein, suppress aspects of the *max2* phenotype that are associated with karrikin responses, but not those related to SL responses, supporting the idea of separate target proteins downstream of MAX2 for KL and SL signalling (Stanga et al, 2013; Soundappan et al, 2015). Several proteins have been suggested as proteolytic targets of SCF^MAX2^ in response to SL signalling, based on biochemical or genetic approaches. One study identified the growth-restricting DELLA transcriptional regulators as targets of SL signalling in rice (Nakamura et al, 2013), while the brassinosteroid response factor BRI1 EMS SUPPRESOR1 (BES1) has been suggested as a candidate in Arabidopsis (Wang et al, 2013). Further studies in rice have identified DWARF53 as a plausible direct target of SCF^MAX2^, since dominant *d53* mutants phenocopy SL resistant mutants, and the D53 protein is degraded in response to treatment with the SL analogue *rac*-GR24 (Zhou et al, 2013; Jiang et al, 2013). Remarkably, D53 is a homologue of SMAX1, suggesting that as with KAI2 and D14, different members of the same protein family mediate separable SL and KL signalling activities. Recent studies in Arabidopsis have shown that the co-orthologues of D53, SMAX1-LIKE6 (SMXL6), SMXL7 and SMXL8, have conserved roles as SL targets in the regulation of development (Soundappan et al, 2015; Wang et al, 2015; Liang et al, 2016). This suggests the attractive hypothesis that the SL signalling pathway evolved through duplication and diversification of proteins both upstream and downstream of MAX2.

Further downstream, most work has focused on the role of SLs in regulating the activity of axillary buds. SL deficient mutants have a highly branched phenotype, leading to the hypothesis that SLs function as negative regulators of shoot branching. In this context the BRANCHED1 (BRC1) TCP-domain transcription factor has been implicated as a transcriptional target of SL, since *brc1-2* mutants have increased, SL-resistant shoot branching (Aguilar-Martinez et al, 2007), and SL can up-regulate *BRC1* expression in pea (Braun et al, 2012). However, this linear model cannot explain the promotion of branching by exogenous SL treatment in genetic backgrounds with compromised auxin transport (Shinohara et al, 2013). This ability of SLs to have both positive and negative effects on branching can be explained by a model in which the PIN1 auxin efflux carrier is a primary downstream target of SL signalling. Consistent with this idea, SL synthesis mutants have increased auxin transport and PIN1 accumulation (Bennett et al, 2006), and *rac*-GR24 can rapidly induce depletion of PIN1 from the plasma membrane of stem xylem parenchyma cells (Shinohara et al, 2013; Crawford et al, 2010).

To clarify the roles of these various proposed SL signalling components and targets in shoot branching control, we have prioritised morphological phenotypic characterisation in relevant genetic backgrounds, which has been less emphasised in some previous studies (Bennett & Leyser, 2014). These analyses are complicated, since that shoot branching is regulated by many factors, the strigolactone analog *rac*-GR24 does not specifically activate the SL signalling pathway (Scaffidi et al, 2013; Scaffidi et al, 2014), and most of the relevant mutants have pleiotropic phenotypes. To overcome these problems, we have used a range of assays for shoot branching, and assessed additional adult shoot phenotypes. Using SL synthesis mutants, we have defined a phenotypic syndrome for the effects of SLs in adult shoot development, and used this to test the role of candidate factors in SL signalling. We show that all the assessed effects of SL in Arabidopsis shoots are mediated through MAX2 and D14, and not the D14 homologues KAI2 or DLK2. We show that mutations in *kai2* do cause some MAX2-dependent phenotypic effects in adult shoots, and that the *max2* adult shoot phenotype is equivalent to a *d14 kai2* double mutant. We demonstrate that BES1 and DELLA proteins are not targets of SL signalling in the regulation of shoot branching, nor likely any other aspect of shoot development. In contrast, we provide further evidence that proteins in the SMXL6/SMXL7 clade are the targets of SL signalling in all the assessed shoot responses, whereas BRC1 and PIN1 are plausible downstream targets of SL signalling specifically in the context of shoot branching, with BRC1 likely acting in parallel to PIN1.

## RESULTS

### Strigolactone influences multiple shoot phenotypes

The most intensively studied aspect of SL developmental responses has been shoot branching, but the phenotypes of SL synthesis mutants include other aspects of adult shoot development. For example, in Arabidopsis SL has been implicated in the control of leaf blade and petiole length, leaf senescence, internode elongation and final height, branch angle, stem diameter, and cambial development (Smith & Waters, 2012; Liang et al, 2016). To provide a baseline for dissecting SL signalling in the adult shoot, we quantified phenotypes in the strong strigolactone synthesis mutant *max4-5* (Bennett et al, 2006). Under our growth conditions, relative to Col-0 wild-type, *max4-5* has greatly increased shoot branching, narrower branch angle, reduced height, reduced stem thickness and delayed leaf senescence (Figure 1B,C,E,F; Figure S1B-C). It also has shorter petioles and leaf blades, but no reduction in blade width, leading to an altered leaf shape (Figure 1A,D; Figure S1A).

**Figure 1:**
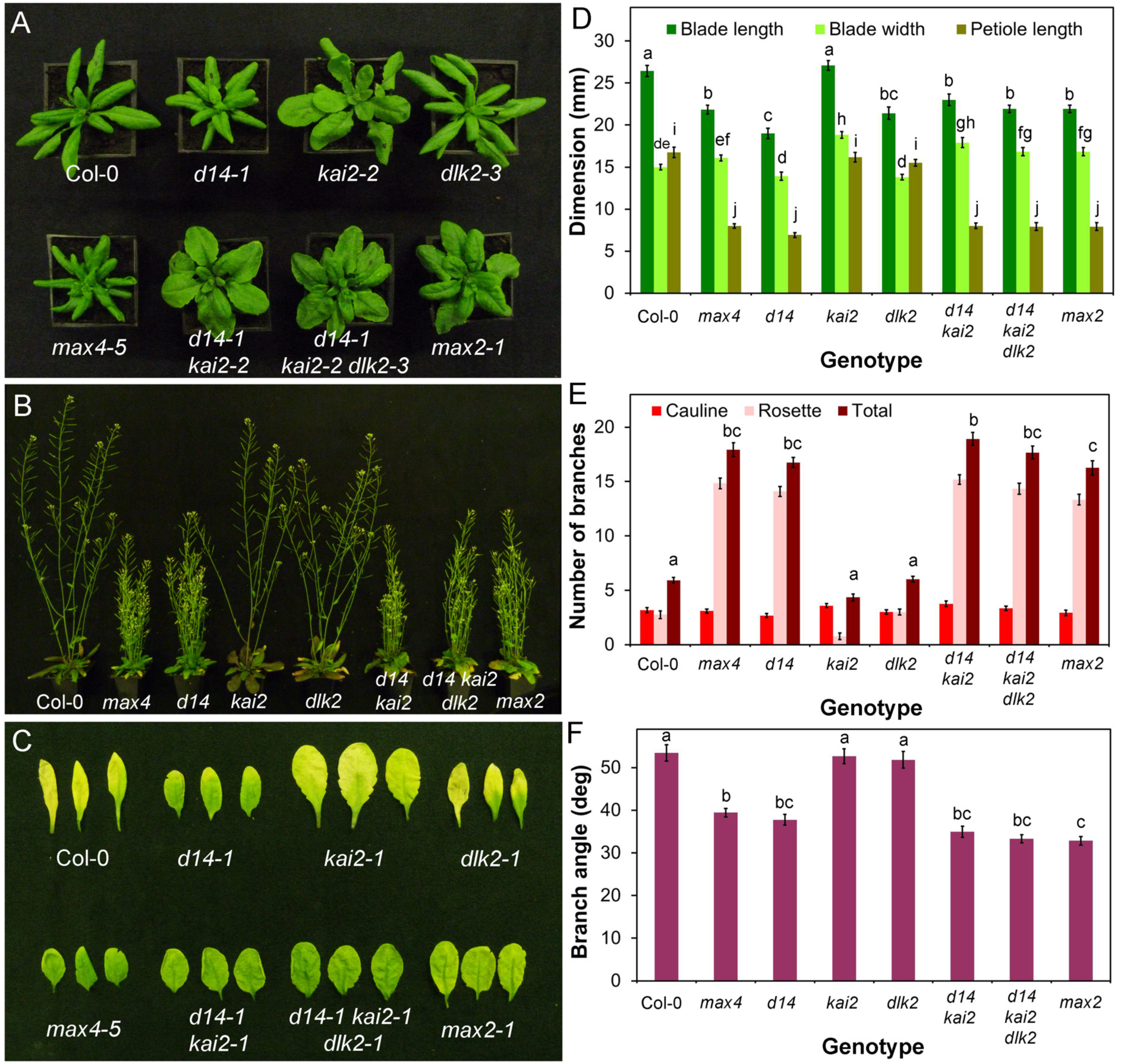
D14 mediates SL signalling in the adult shoot. **A)** Rosette leaf phenotypes in candidate SL signalling mutants 4 weeks after germination. **B)** Branching phenotypes in candidate SL signalling mutants 6 weeks after germination. **C)** Dark-induced leaf senescence phenotypes in candidate SL signalling mutants. Rosette leaves were wrapped in foil for 8 days then imaged. **D)** Leaf dimensions in candidate SL signalling mutants. Measurements were made on the 7^th^ rosette leaf, 35 days after germination. n=10-12, bars indicate s.e.m. Bars with the same letter are not significantly different from each other (ANOVA, Tukey HSD test). **E)** Branching levels in candidate SL signalling mutants. Numbers of primary cauline and rosette branches were measured at proliferative arrest, n=10-12, bars indicate s.e.m. Bars with the same letter are not significantly different from each other (ANOVA, Tukey HSD test). **F)** Branch angle (measured in degrees) in candidate SL signalling mutants,, n=10-12, bars indicate s.e.m. Bars with the same letter are not significantly different from each other (ANOVA, Tukey HSD test).

Having established a phenotypic platform for understanding the effects of SL deficiency in adult shoots, we tested whether mutations in proposed or potential SL signalling genes confer the expected phenotypic profile. For positive regulators of SL signalling, loss-of-function, hypomorphic mutants should phenocopy the *max4-5* phenotype, and gain-of-function, hypermorphic mutants should suppress the phenotype of SL deficient/insensitive mutants. For negative regulators these expectations are inverted. Mutants in downstream effectors should display changes in the SL-sensitivity of relevant phenotypes. In practice, the genetic materials do not exist to assess all these aspects for each candidate gene, and genetic analysis is often complicated by problems of pleiotropy, redundancy and epistasis. Nevertheless, we were able to gather sufficient materials for each candidate to assess their role in SL signalling.

### SL signalling in the Arabdopsis adult shoot is mediated by D14

As discussed above, two proteins are known to be required for SL signalling, MAX2 and D14. The leaf dimensions and leaf senescence, branching level, branch angle, height and stem thickness phenotypes of *d14-1* are essentially indistinguishable from *max4-5* (Figure 1; Figure S1). Consistent with previous reports (e.g.Waters et al, 2012a; Chevalier et al, 2014), we also found that *d14-1* is strongly SL insensitive in a branching assay (t-test, n=12, p=0.179) (Figure S1D).. By contrast, we did not observe any clear phenocopy of *max4-5* in the *kai2-2* or *dlk2-3* mutants (Figure 1; Figure S1). The *kai2* mutant has distinct phenotypic effects in the shoot that are not seen in *max4-5*, including strongly accelerated flowering time (Figure S1E) and increased leaf blade width (Figure 1A). In contrast, the *dlk2-3* mutant is largely indistinguishable from wild-type, though there are subtle effects in leaf size and height in this line (Figure 1, Figure S1). We conclude that D14-dependent signalling is fully responsible for SL effects on shoot branching.

In contrast to *d14-1*, the *max2-1* mutant is not a simple phenocopy of *max4-5* (Figure 1A). Most aspects of the *max4-5* adult shoot phenotype are evident within the *max2-1* phenotype, including increased shoot branching, reduced height, decreased petiole length and delayed leaf senescence (Figure 1; Figure S1A). However, there are additional phenotypes, including wider leaf blades. Since MAX2 has been implicated in signalling downstream of KAI2, we reasoned that the *max2-1* phenotype may represent combined loss of function of signaling downstream of these two receptors, which we confirmed by making a *d14-1 kai2-2* double mutant, which closely phenocopies *max2-1* (Figure 1). This interaction most clearly illustrated by leaf shape (Figure 1A and D), which combines characteristics of the single mutants to produce *max2*-like leaves.

We reasoned that if DLK2 acted redundantly with D14 or KAI2, the effect of losing DLK2 would be more obvious in the sensitized *d14-1 kai2-2* background. We thus examined a *d14-1 kai2-2 dlk2-3* mutant, but did not observe any clear evidence of enhancement of phenotypes relative to *d14-1 kai2-2* (Figure 1, Figure S1). Given the similarity of the *d14-1* and *max4-5* phenotypes, and the lack of obvious redundancy with KAI2 and DLK2, we conclude that for all the phenotypes we examined, SL signalling is mediated by D14 acting through MAX2.

### DELLA proteins are not targets of SL signalling in shoot branching

We next assessed whether proteins that have been previously implicated as direct proteolytic targets of SCF^MAX2^ show the expected phenotypes of negatively regulated targets. We first examined the DELLA proteins, constitutive repressors of growth that are degraded in the presence of gibberellins (GA). DELLA proteins have been identified as SL signalling targets based on their physical interactions with D14 (Nakamura et al, 2013). We used the dominant negative *gai* mutant in which the GAI DELLA protein is stabilized, phenocopying severe GA deficiency (Peng et al, 1997), and the quintuple *gai-t6 rga-t2 rgl1-1 rgl2-1 rgl3-1* (‘*della*’) mutant, in which all DELLA protein activity is lost (Feng et al, 2008). These mutations confer extreme and opposite changes in growth habit. The *gai* mutant is dwarfed, with short leaves and internodes, and grows slowly, while *della* has long internodes, long leaves and develops at an increased rate, flowering early (Figure 2A-C). We assessed whether these mutants have any phenotypic overlap with SL synthesis or signalling mutants. There are clear leaf phenotypes in both *gai* and *della* mutants (Figure 2D), but these do not alter the relative shape of the leaf (length/width ratio) (ANOVA, Tukey HSD, n=9−10, p>0.05), only the absolute dimensions of the leaf (Figure S2A,B). The effect of DELLA activity on leaves is thus qualitatively different from the effect of SL signalling. There was also no alteration in leaf senescence in *della* relative to L*er*, but there may be a delay in the *gai-1* mutant (Figure S2C). As anticipated, height was increased in *della*, and reduced in *gai* relative to L*er* (Figure S2D). With respect to height, the effect of *gai* is thus qualitatively similar to SL mutants, but is quantitatively much more extreme. Stem diameter follows the same pattern, being increased in *della*, and reduced in *gai* (Figure S2E). We observed no difference in branch angle between L*er* and *gai*, but branch angle was increased in *della* (Figure S2F). Finally, we examined whether either mutant had a branching phenotype under standard long-day growth conditions, but did not observe any statistical difference from the L*er* wild-type in terms of total primary branches in *della* or *gai* (ANOVA, Tukey HSD test, n=13−20, p>0.05)(Figure 2H). The distribution of branches between cauline and rosette nodes was altered (Figure 2H), but this is attributable to differences in the number of cauline nodes produced in *gai*/*della*. We also trialled a more sensitive decapitation-based assay to assess branching (Greb et al, 2003), but found that this was unsuitable in the L*er* background, due to precocious outgrowth of rosette buds before decapitation, which does not normally occur in Col-0.

**Figure 2:**
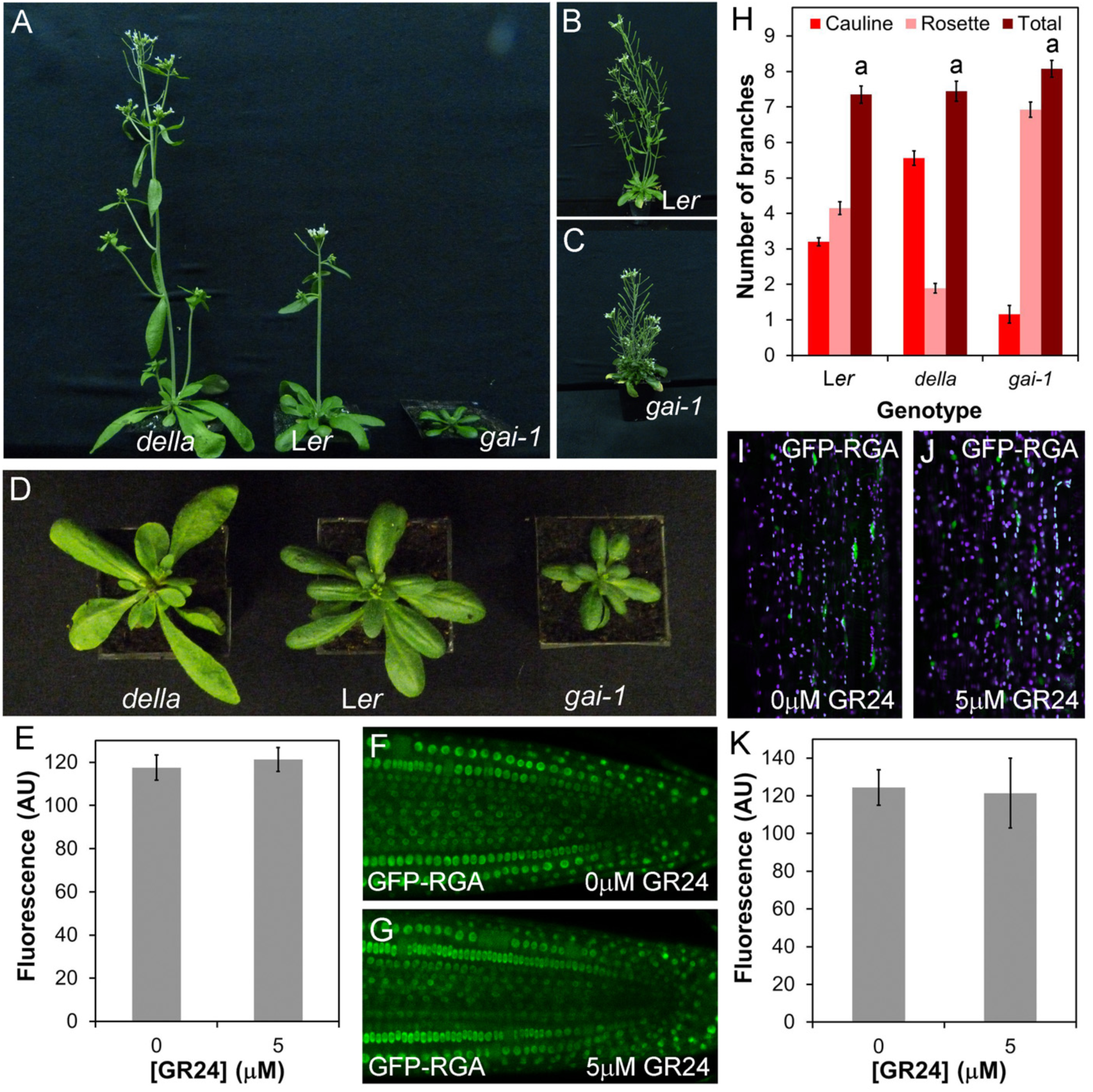
DELLA proteins are not targets of SL signalling in shoot development. **A)** Shoot morpohology in age-matched plants of *gai-t6 rga-t2 rgl1-1 rgl2-1 rgl3-1 (della), Ler and gai-1*. **B)** *Ler* plant at later developmental stage than A) showing branching habit. **C)** *gai-1* plant at later developmental stage than A) showing branching habit. **D)** Rosette morphology phenotypes in age-matched plants of *della*, Ler and *gai-1*. **E-G**) Effect of *rac*-GR24 treatment on stability of the GFP-RGA fusion protein in roots. F) and G) show representative images of roots treated with 0μM or 5μM *rac*-GR24 for 45 minutes respectively, and E) shows quantification of relative fluorescence in the two treatments; n=5 nuclei in each of 12 roots per treatment. The mean value per root is shown, along with the standard error of this mean. **H)** Numbers of primary branches in long-day grown *Ler*, *della* and *gai-1* plants, measured at proliferative arrest, n=13-20, bars indicate s.e.m. Bars with the same letter are not significantly different from each other (ANOVA, Tukey’s HSD test). **I-K)** Effect of *rac*-GR24 treatment on stability of the GFP-RGA fusion protein in shoots. J) and K) show representative images of hand sectioned 6-week old stems treated with 0μM or 5μM *rac*-GR24 for 45 minutes respectively, and I) shows quantification of relative fluorescence in the two treatments; n=5 nuclei in each of 8 shoots per treatment. The mean value per stem is shown, along with the standard error of this mean.

From our phenotypic analysis, although the *gai* and *della* mutants share some phenotypic characteristics with reduced and increased SL signalling mutants respectively, their phenotypic syndromes and the correlations within them are both qualitatively and quantitatively different. It is therefore plausible, if unlikely, that SL could regulate some aspects of shoot phenotype by targeting DELLA proteins for degradation. To assess more directly the effects of SL on DELLA stability, we treated roots expressing a GFP-RGA fusion protein (Fu et al, 2003) with 5µM *rac*-GR24 for 45 minutes (a relevant timeframe for SL action), but observed no decrease in the level of fluorescence of the fusion protein relative to mock-treated plants (t-test, n=12, p=0.645)(Figure 2F-H). We then repeated this analysis in hand-sectioned, 6-week old primary inflorescence stems, but again, found no effect of *rac*-GR24 on RGA stability (Figure 2I-K)(t-test, n=8, p=0.88). We thus conclude that SL is unlikely to control development through direct effects on DELLA proteins. Consistent with this idea, SL acts independently of GA and DELLAs in the control of internode elongation in pea (de Saint Germain et al, 2013b).

### BES1 is not a target of SL signalling in shoot branching

BES1, a transcription factor which regulates brassinosteroid (BR) responses along with its homologues BZR1 and BEH1-BEH4, has been proposed as a direct target of SL signalling based primarily on biochemical approaches (Wang et al, 2013). Consistent with this idea, the gain of function *bes1-D* mutant (in which BES1 is stabilized) was reported to have increased branching, while *BES1*-*RNAi* lines were reported to have reduced branching (Wang et al, 2013). However, no other BR-related mutants have been reported to have branching phenotypes, and BR has not previously been implicated in the regulation of branching, or in SL responses. We thus re-examined the role of BES1 in shoot branching. We obtained the original *bes1-D* line (Yin et al, 2002) from the Nottingham Arabidopsis Stock Centre (NASC), and found that the line contains multiple segregating phenotypes, including increased shoot branching, but this phenotype does not appear to be linked to the characteristic *bes1-D* leaf phenotype, suggesting that the branching defect reported by Wang et al may be wrongly attributed to mutation in BES1. In order to circumvent these issues, we obtained and characterized a verified *bes1-D* line that had been backcrossed multiple times to the Col-0 wild-type (Gonzalez-Garcia et al, 2011), as well as a loss-of-function T-DNA allele, *bes1-1* (He et al, 2005) The *bes1-D* mutant has a characteristic leaf phenotype (Figure 3A), but this is qualitatively different from the SL mutant leaf phenotype and results from increased blade width as well as uneven lamina expansion. Petiole and blade length are not significantly different from wild-type (ANOVA, Tukey HSD, n=9-10, p>0.05). There is no difference in any leaf dimension between *bes1-1* and Col-0 (Figure S3A)(ANOVA, Tukey HSD, n=9-10, p>0.05), and leaf senescence is not delayed in *bes1-D* or *bes1-1* relative to Col-0 (Figure S3B). We observed no significant difference in height between Col-0, *bes1-1* and *bes1-D* (ANOVA, Tukey HSD, n=10, p>0.05)(Figure S3C), and no difference in stem diameter between Col-0 and *bes1-1*, though there is a significant reduction in *bes1-D* relative to Col-0 (ANOVA, Tukey HSD, n=10, p<0.05) (Figure S3D)..There is also a significant increase in branch angle in *bes1-D* relative to Col-0, but branch angle in *bes1-1* is not different from Col-0 (Figure S3E) (ANOVA, Tukey HSD, n=10, p>0.05).

**Figure 3:**
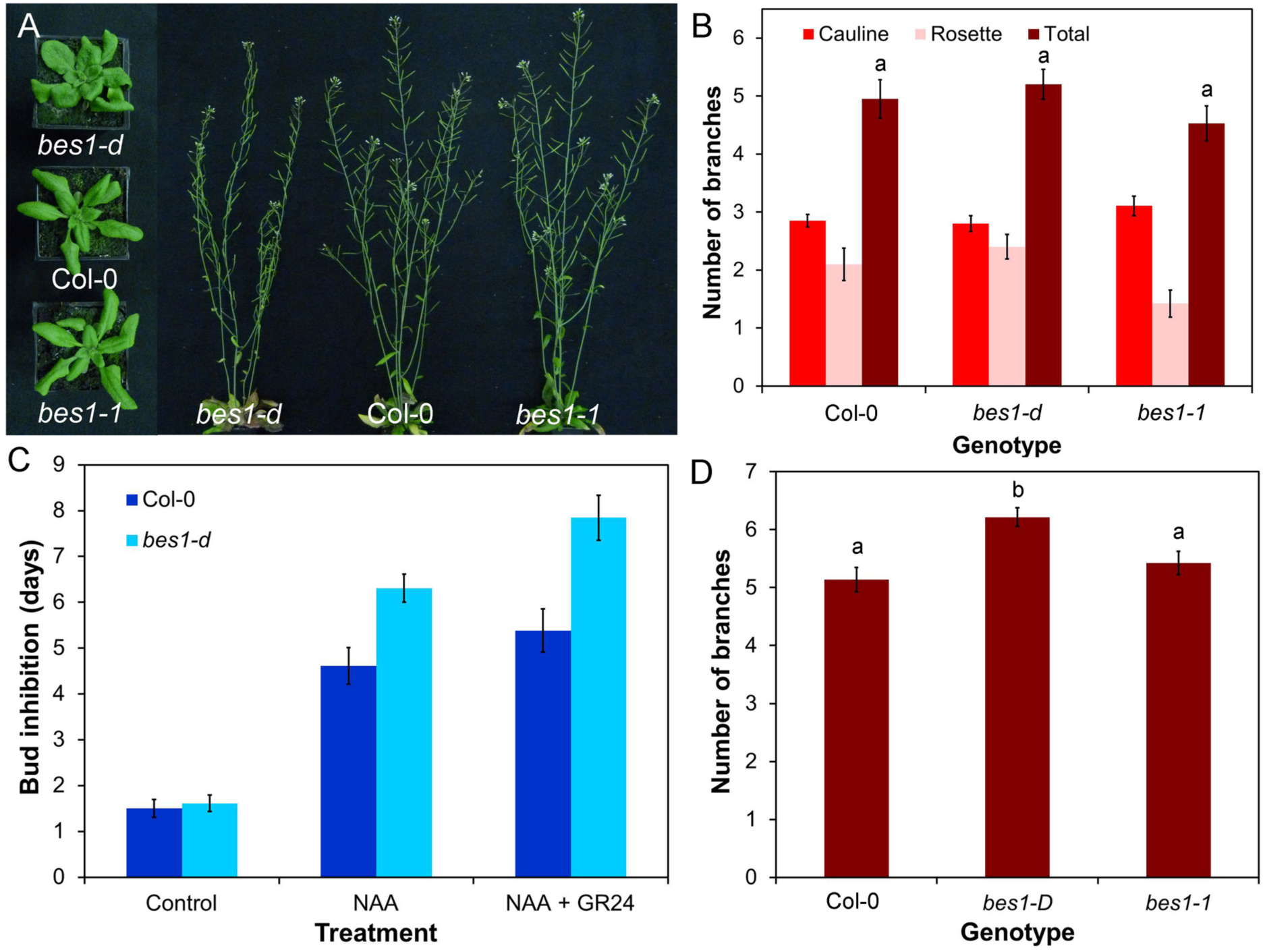
BES1 is not a target of SL signalling in shoot branching. **A)** Leaf and branching phenotypes in Col-0*, bes1-D* and *bes1-1* at 4 and 6 weeks post germination respectively. **B)** Numbers of primary branches in long-day grown Col-0, *bes1-D* and *bes1-1*. Branching was measured at proliferative arrest, n=19-20, bars indicate s.e.m. Bars with the same letter are not significantly different from each other (ANOVA, Tukey HSD test). **C)** Growth responses of Col-0 and *bes1-D* buds on excised nodal stem segments. Stem segments were treated with either solvent control, 1µM NAA applied apically, or 1µM NAA apically + 5µM *rac*-GR24 basally. The mean number of days that buds took to reach a length greater than 1.5mm is shown for each genotype and treatment, n=12-13 nodes per treatment, bars indicate s.e.m. **D)** Numbers of primary rosette branches in decapitated Col-0, *bes1-D* and *bes1-1* plants grown in short photoperiods and then shifted to long photoperiods, 10 days after decapitation. n=22-37, bars indicate s.e.m. Bars with the same letter are not significantly different from each other (ANOVA, Tukey HSD test).

We found that neither *bes1-D* nor *bes1-1* show any difference in branching levels relative to Col-0 in a standard long day assay (ANOVA, Tukey HSD, n=20, p>0.05) although *bes1-d* (but not *bes1-1*) shows a slight increase in branching in the more sensitive decapitation-based assay [41](ANOVA, Tukey HSD, n=22-37, p<0.05)(Figure 3A,B,D). We also tested whether knocking out *BES1* reduces branching in a *max2-1* background, but found that the *bes1-1 max2-1* double mutant produces the same number of branches as *max2-1* (ANOVA, Tukey HSD, n=19-20, p>0.05)(Figure S3F). This result contrasts to previous reports that *BES1*-*RNAi* lines suppress the branching phenotype of *max2-1.* The *BES1*-*RNAi* lines have highly pleiotropic phenotypes and are generally lacking in vigour, making the results difficult to interpret (Wang et al, 2013).

It has also been suggested that *bes1-D* alters sensitivity to SL, because the SL analog *rac*-GR24 does not reduce hypocotyl length in the *bes1-D* background (Wang et al, 2013). We therefore tested whether *bes1-D* axillary buds are insensitive to *rac*-GR24, using a well-established excised node assay. In this assay, *rac*-GR24 treatment can enhance the inhibitory effects of apically applied auxin on bud growth. We found that *bes1-D* is fully sensitive to *rac*-GR24 in this assay (t-test, n=13, p<0.01). Indeed the kinetics of bud outgrowth in response to either NAA or NAA + *rac*-GR24 treatment are slightly retarded relative to wild-type, rather than accelerated as would be predicted if BES1 is a target for SL signalling in this response (Figure 3C). Thus the *bes1-D* mutation neither increases shoot branching, nor reduces bud SL responses.

### SMXL6 is functionally similar to SMXL7

Recent analysis of SMXL6, SMXL7 and SMXL8 has suggested that they are major targets of SL signalling in Arabidopsis (Soundappan et al, 2015; Wang et al, 2015; Liang et al, 2016). Combined loss-of-function of these three genes is sufficient to suppress the branching, height, leaf/petiole length and lateral root density phenotypes of *max2* that are associated with SL signalling deficiency, but does not affect the germination, hypocotyl length or leaf width phenotypes of *max2* that are associated with KAI2-mediated signalling (Soundappan et al, 2015). Based on these loss-of-function phenotypes, it is clear that in Arabidopsis, SMXL7 plays the dominant role (Soundappan et al, 2015), and as such has received more attention (Liang et al, 2016). We have recently shown that expression of stabilized SMXL7 is sufficient to recapitulate all examined aspects of the SL phenotypic syndrome (Liang et al, 2016). An interesting question is whether SMXL6 and SMXL8 demonstrate similar behaviour and functionality, despite their subordinate role in regulating development. It is for instance possible that SMXL6 and SMXL8 actually have rather different functions to SMXL7, and only act in a SMXL7-like manner in the absence of that protein, e.g. analogous to APETALA1, CAULIFLOWER and FRUITFULL in the control of shoot meristem fate (Ferrandiz et al, 2000).

To assess the behavior of SMXL6, we created a SMXL6-YFP fusion, expressed from the 35S promoter (*35Spro:SMXL6-YFP*), and transformed it into Arabidopsis. As with SMXL7, we observed a clear nuclear localization for SMXL6 in cells of the Arabidopsis root meristem (Figure 4B). Similar to SMXL7, we struggled to detect SMXL6-YFP in wild-types stems, but in the stabilizing *max2-1* background, we detected SMXL6-YFP in the nucleus of vascular-associated cells (Figure 4A). We tested whether SMXL6 also shows the rapid *rac*-GR24 induced degradation we observed for SMXL7, and found that SMXL6 protein levels are greatly reduced in the root meristem after 20 minutes treatment with 5µM *rac*-GR24 (Figure 4B-F), thus displaying very similar kinetics to SMXL7 (REFs). This response was blocked in a *max2-1* background or in the presence of the 26S proteasome inhibitor MG132 (Figure 4H,I,J), and did not occur in response to treatment with 1µM KAR1 (a karrikin) (Figure 4G). We also created a version of SMXL6 lacking the ‘p-loop’ required for SCF^MAX2^-mediated degradation (Zhou et al, 2013; Jiang et al, 2013; Soundappan et al, 2015), and then expressed this under the 35S promoter in the Col-0 background (*35S:SMXL6*^Δ^*pl*^^-*YF*P). As anticipated, SMXL6^Δ^*pl*^^-YFP was resistant to *rac*-GR24 induced degradation (Figure 4K,L). We thus conclude that the general behavior of SMXL6 is very similar to that described for SMXL7 [Liang et al, 2016].

**Figure 4:**
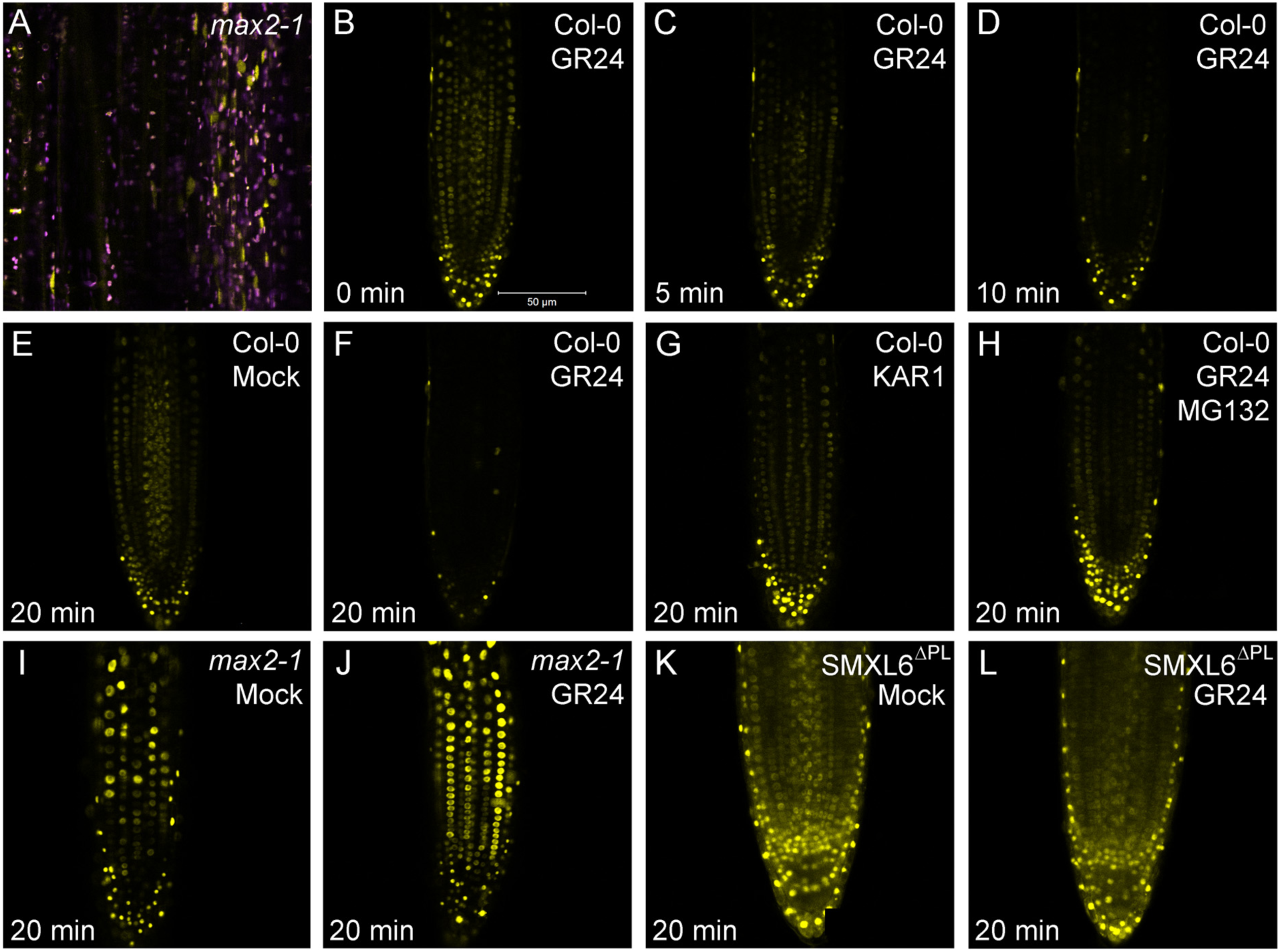
SMXL6 is degraded in response to SL treatment **A)** Expression of SMXL6-YFP in vascular cambium cells of *max2-1* stems (yellow). Purple signal indicates chloroplast autofluorescence. **B-D)** Response of SMXL6-YFP protein levels in Col-0 roots to treatment with 5µM *rac*-GR24 over a 10 minute time course. **E-H)** Comparison of SMXL6-YFP protein levels in Col-0 roots after 20 minutes treatment with solvent control (E) 5µM KAR1 (G) or 5µM *rac*-GR24 in the presence (H) or absence (F) of MG132, an inhibitor of the 26S proteasome. **I,J)** Comparison of SMXL6 protein levels in *max2-1* roots after 20 minutes treatment with solvent control (I) or 5µM *rac*-GR24 (J). **K,L)** Comparison of SMXL6^Δ^pl^^-YFP protein levels in roots after 20 minutes treatment with solvent control (K) or 5µM *rac*-GR24 (L)

We next assessed the developmental potential of the SMXL6 protein using *35S: SMXL6*^Δ^*pl*^^-*YFP* transgenic lines.. We observed that multiple independent stably transformed lines had a phenotype closely resembling that of SL deficient mutants (also observed in Wang et al, 2015). We quantified shoot phenotypes in a representative line (Figure 5). In terms of shoot branching, *35S:SMXL6*^Δ^*pl*^^-*YFP* confers similar phenotypes to those seen in *d14-1* and *max2-1*, if somewhat less extreme (ANOVA, Tukey HSD test, n=10-12, p<0.05)(Figure 5C,E); there is a similar effect on final height (Figure S4A) The buds of *35S:SMXL6*^Δ^*pl*^^-*YFP* plants are insensitive to the application of *rac*-GR24 when tested in an excised node assay (t-test, n=13, p=0.39) (Figure 5E,F). The leaf phenotype of *35S:SMXL6*^Δ^*pl*^^-*YFP* is intermediate between *d14-1* and *max2-1*, with the characteristic short petioles of SL mutants (Figure 5A,D). *35S:SMXL6*^Δ^*pl*^^-*YFP* leaves are slightly wider and shorter than wild-type (ANOVA, Tukey HSD, n=11-12, p<0.05). They have the same blade length:width ratio as *max2-1* (ANOVA, Tukey HSD, n=11-12, p<0.05)(Figure S4B), but are not as large as *max2-1* leaves (Figure 5D). Whilst we intuitively expected *35S:SMXL6*^Δ^*pl*^^-*YFP* leaves to resemble *d14-1* rather than *max2*, very similar *max2*-like phenotypes were also observed in lines expressing SMXL7 from the *35S* promoter [Liang et al, 2016]. This *max2*-like phenotype suggests that the use of the 35S promoter produces some off-target effects, for example on KAI2-related signalling We also tested the involvement of SMXL6 and SMXL7 in leaf senescence, which has not previously been assessed. We found that like *d14-1* and *max2-1*, *35S:SMXL6*^Δ^*pl*^^-*YFP* causes delayed senescence in leaves placed in the dark for 7 days (Figure 5B). Conversely, we found that loss-of-function mutation of SMXL6 and SMXL7 was sufficient to suppress the *max2-1* leaf senescence phenotype (Figure 5B). Thus SMXL6 and homologous proteins also contribute to dark-induced leaf senescence.

**Figure 5:**
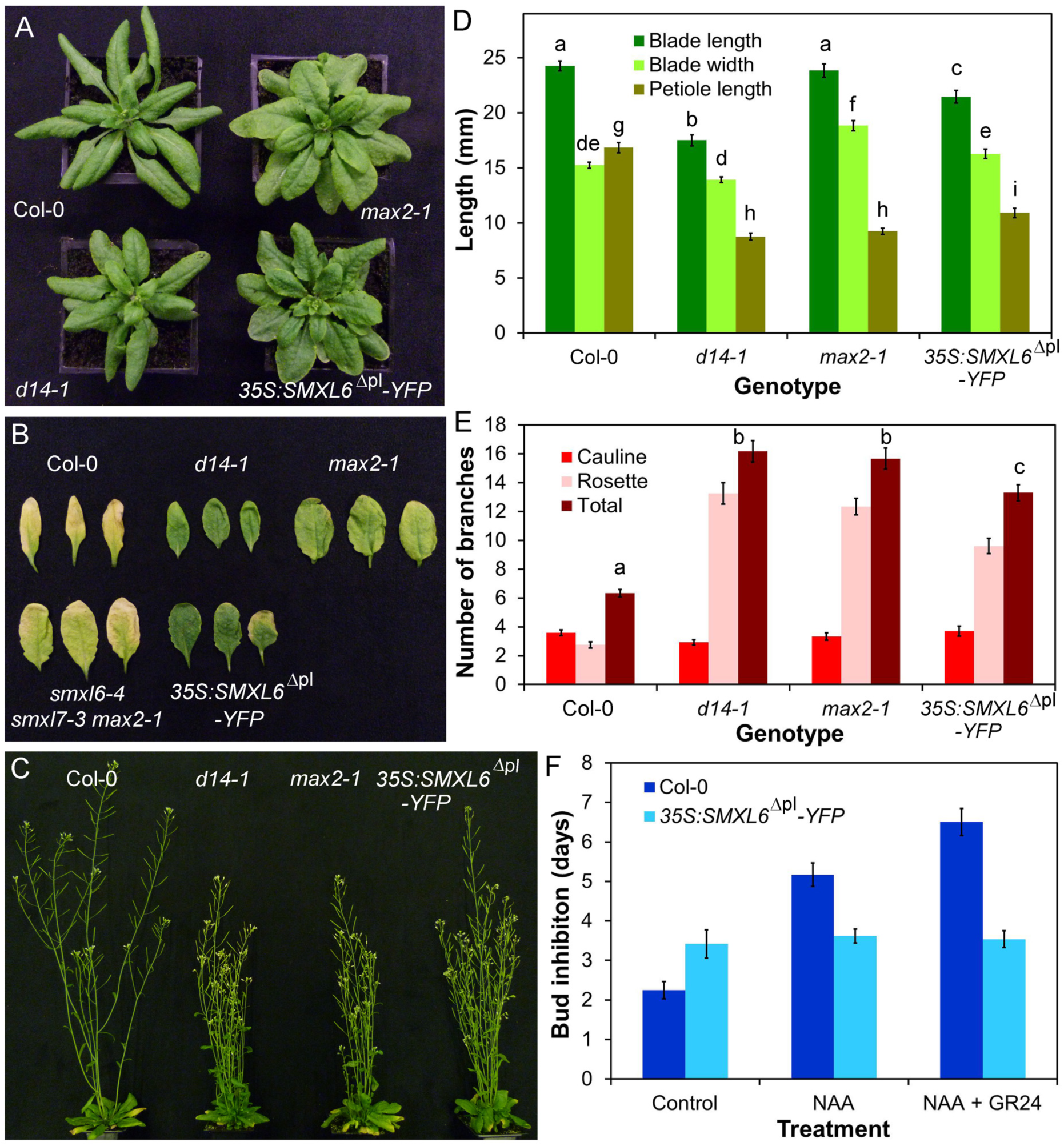
SMXL6 is functionally similar to SMXL7. **A)** Rosette leaf phenotypes in 4 week old Col-0, *max2-1*, *d14-1*, and *35S:SMXL76*^Δ^*pl*^^-*YFP* plants. **B)** Dark-induced senescence in Col-0*, d14-1*, *max2-1*, *smxl6-4 smxl7-1 max2-1* and *35S:SMXL76*^Δ^*pl*^^-*YFP* leaves from 5 week old plants. Leaves were wrapped in foil and imaged after 7 days. **C)** Branching phenotypes in 6 week old Col-0*, d14-1*, *max2-1* and *35S:SMXL76*^Δ^*pl*^^-*YFP* plants. **D)** Leaf dimensions in Col-0, *d14-1*, *max2-1* and *35S:SMXL76*^Δ^*pl*^^-*YFP* lines. Measurements were made on the 7^th^ rosette leaf, 35 days after germination. n=11-12, bars indicate S.E.M. Bars with the same letter are not significantly different from each other (ANOVA, Tukey HSD test). **E)** Numbers of primary rosette branches in long-day grown Col-0*, d14-1*, *max2-1* and *35S:SMXL76*^Δ^*pl*^^-*YFP.* Number of primary rosette branches was measured at proliferative arrest, n=10-12, bars indicate S.E.M. Bars with the same letter are not significantly different from each other (ANOVA, Tukey HSD test). **F)** Growth responses of Col-0 and *35S:SMXL76*^Δ^*pl*^^-*YFP* buds on excised nodal sections. Nodes were treated with either solvent control, 0.3µM NAA applied apically, or 0.3µM NAA apically + 5µM *rac*-GR24 basally. The mean number of days that buds took to reach a length greater than 2mm is shown for each genotype and treatment, n=12-16 nodes per treatment, bars indicate s.e.m

### BRC1 and BRC2 regulate shoot branching and stature

We next examined the role of putative downstream targets in SL responses. *BRC1* has been suggested as a transcriptional target of SL signalling, based on the SL-resistant increased shoot branching phenotype observed in *brc1* loss of function mutants, and the lack of genetic additivity in some, but not all, *brc1 max* double mutants (Aguilar-Martinez et al, 2007; Braun et al, 2012; Chevalier et al, 2014). We assessed whether *BRC1* could be a more general target of SL response. Consistent with previous reports, we observed a large increase in rosette branching in *brc1-2 brc2-1*, although in our conditions less so than in *max4-5*, *d14-1* and *max2-1* (ANOVA, Tukey HSD, n=12, P<0.05)(Figure 6B,C)..We found that flowering is accelerated in *brc1-2 brc2-1* relative to Col-0 (t-test, n=11-12, p<0.005)(Figure S5A). The resultant reduction in leaf number, and hence axillary bud number, could account for some of differences in branching relative to *max4-5*. In addition, the early flowering of axillary shoots could account at least in part for the increased number of elongated branches compared to wild-type (Niwa et al, 2013). We found no clear effect of *brc1-2 brc2-1* on blade length, blade width, petiole length, leaf shape or leaf senescence (Figure 6A,D; Figure S5B). However, plant height is reduced in *brc1-2 brc2-1*, although not to the same extent as seen in *d14-1* (ANOVA, Tukey HSD, n=12, p<0.05)(Figure S5C). These data suggest that *BRC1* is a plausible target of SL signalling, although only in the contexts of shoot branching and stature. However, they also show that, for these responses, loss of *BRC1* and *BRC2* expression cannot explain the full phenotypic effect of deficient SL signalling.

**Figure 6:**
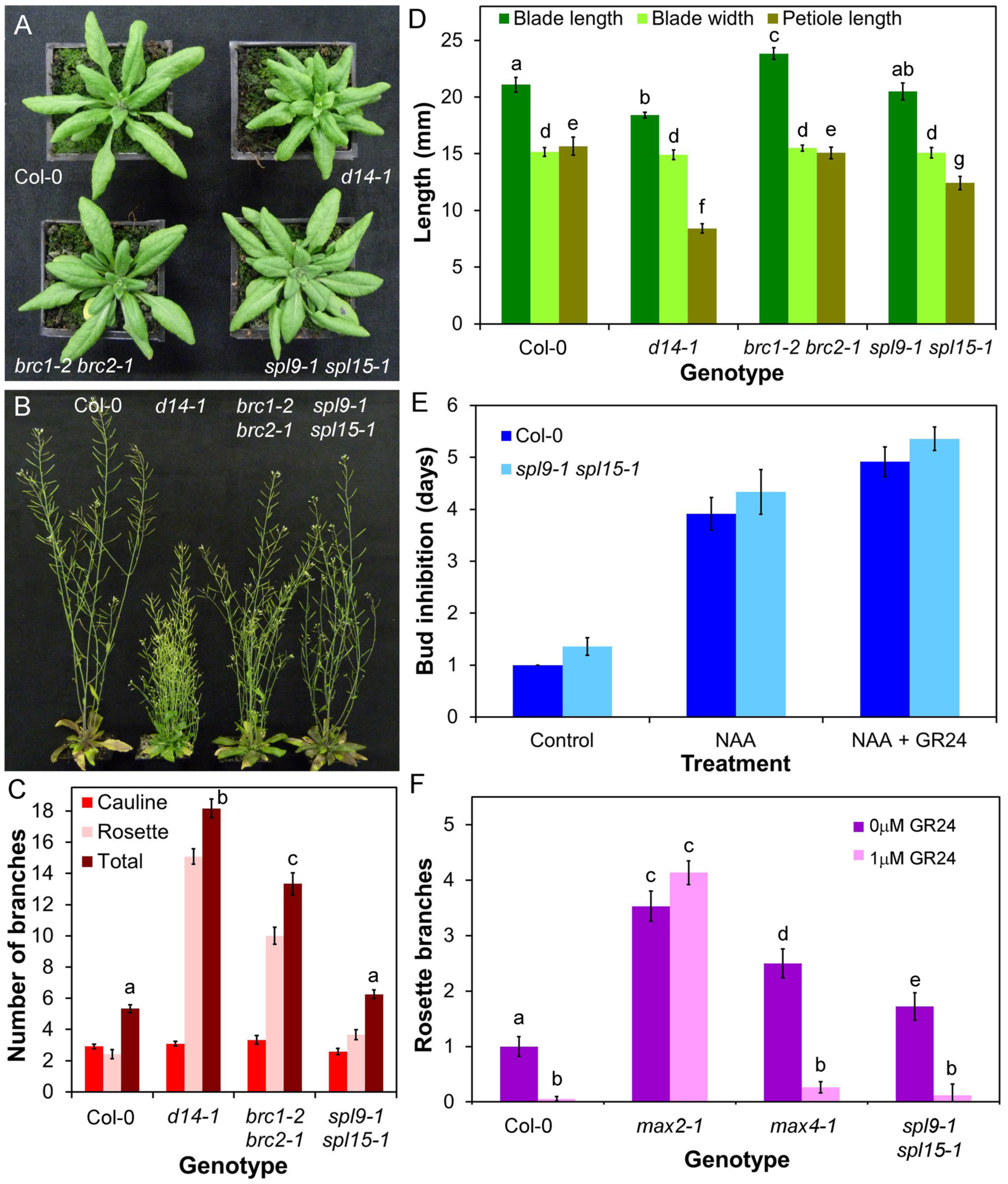
The role of BRC1/BRC2 and SPL9/SPL15 in shoot development. **A)** Rosette leaf phenotypes in 4 week old Col-0*, d14-1*, *brc1-2 brc2-1* and *spl9-1 spl15-1* plants. **B)** Branching phenotypes in 6 week old Col-0*, d14-1*, *brc1-2 brc2-1* and *spl9-1 spl15-1* plants. **C)** Numbers of primary rosette branches in long-day grown Col-0*, d14-1*, *brc1-2 brc2-1* and *spl9-1 spl15-1.* Number of primary rosette branches was measured at proliferative arrest, n=12, bars indicate s.e.m. Bars with the same letter are not significantly different from each other (ANOVA. Tukey HSD test). **D)** Leaf dimensions in candidate SL signalling mutants. Measurements were made on the 7^th^ rosette leaf, 35 days after germination. n=12, bars indicate s.e.m. Bars with the same letter are not significantly different from each other (ANOVA, Tukey HSD test). **E)** Growth responses of Col-0 and *spl9-1 spl15-1* buds on excised nodal sections. Nodes were treated with either solvent control, 0.5µM NAA applied apically, or 0.5µM NAA apically + 5µM *rac*-GR24 basally. The mean number of days that buds took to reach a length greater than 2mm is shown for each genotype and treatment, n=11-14 nodes per treatment, bars indicate s.e.m. **F)** Numbers of primary rosette branches in Col-0*, max2-1*, *max4-1* and *spl9-1 spl15-1* grown on agar solidified media supplemented with 1µM *rac*-GR24 or a solvent control. Number of primary rosette branches was measured at proliferative arrest, n=15-36, bars indicate s.e.m. Bars with the same letter are not significantly different from each other (ANOVA, Tukey HSD test).

### SPL9 and SPL15 are not components of SL response

SPL9 and SPL15 are the closest Arabidopsis relatives of the OsSPL14 gene from rice, which is a negative regulator of shoot branching (Jiao et al, 2010). Both genetic and physical interactions between *OsSPL14* and the rice *BRC1* orthologue have been described, leading to the hypothesis that *BRC1* transcription is regulated by OsSPL14 (Lu et al, 2013). In Arabidopsis, the *spl9-1 spl15-1* double mutant has previously been shown to have increased shoot branching (Schwarz et al, 2008), as have lines overexpressing the micro-RNA miR156, which down-regulates expression of several SPL genes, including *SPL9* and *SPL15* (Schwab et al, 2005; Xing et al, 2010; Wei et al, 2012). A study in rice demonstrated that OsSPL14 acts in a separate pathway to SL signalling (Luo et al, 2012). To investigate the relationship between SL and *SPL9*/*SPL15* we assessed the branching phenotypes of the *spl9-1 spl15-1* double mutant. Under our growth conditions we observed only a very modest increase in branching in *spl9-1 spl15-1*, considerably less than that seen in *d14-1* or *brc1-2 brc2-1* (ANOVA, Tukey HSD, n=12, p<0.05) (Figure 5B,C). We then tested whether, like *brc1-2 brc2-1*, shoot branching in *spl9-1 spl15-1* displays SL resistance. We grew plants on media containing 1µM *rac*-GR24, and observed that this treatment reduced branching in *spl9-1 spl15-*1, to levels similar to wild-type (ANOVA, Tukey HSD, n=15-36, p<0.05) (Figure 6F). We also tested whether *spl9-1 spl15-1* is insensitive to *rac*-GR24 treatment in the excised node assay, and again found that bud outgrowth in these plants is fully sensitive to *rac*-GR24 treatment (t-test, n=12-14, p<0.05) (Figure 6E). We also found that *spl9-1 spl15-1* leaves do not resemble *d14-1* leaves, although they do have a slightly different shape to wild-type leaves (Figure 6A,D). Thus although *spl9-1 spl15-1* mutants do have somewhat increased shoot branching, the phenotypic dissimilarity to *d14-1* and the lack of SL-resistance in the *spl9-1 spl15-1* mutant strongly suggests that SPL9 and SPL15 are not downstream targets of SL signalling, but rather regulate branching through a separate mechanism, as previously suggested in rice (Luo et al, 2012).

### Canonical SL signalling in the shoot modulates auxin transport and PIN1 levels

We have previously shown that the SL synthesis mutants *max1-1*, *max3-9* and *max4-1* have increased auxin transport in the primary inflorescence stem, and that *max1-1* and *max3-9* have increased levels of the PIN1 auxin efflux carrier at the basal plasma membrane of cambial and xylem parenchyma cells in the stem (Bennett et al, 2006). We observed the same effects in *max4-5* and the more recently identified SL synthesis mutant *d27-1* (Figure 7A, Figure 8A-D,I). These phenotypes are also seen in the *max2-1* SL signalling mutant (Figure 7A, Figure 8A,B,I) (Crawford et al, 2010), and we thus tested whether these effects are mediated by *d14-1*, *kai2-1* or *dlk2-1* dependent signalling. We found that auxin transport is increased in the primary inflorescence stems of *d14-1* to the same or greater extent as *max2-1* and *max4-5* (ANOVA, Dunnett’s test, n=18-20, P<0.05), while there is no change in auxin transport in *kai2-1* (here in the Ler background) or *dlk2-1* relative to wild-type (Figure 7A). Similarly, we found that PIN1 levels are increased in *d14-1*, but not *kai2-1* or *dlk2-1* (ANOVA, Tukey HSD, n=8, P<0.05) (Figure 8E-G).

**Figure 7:**
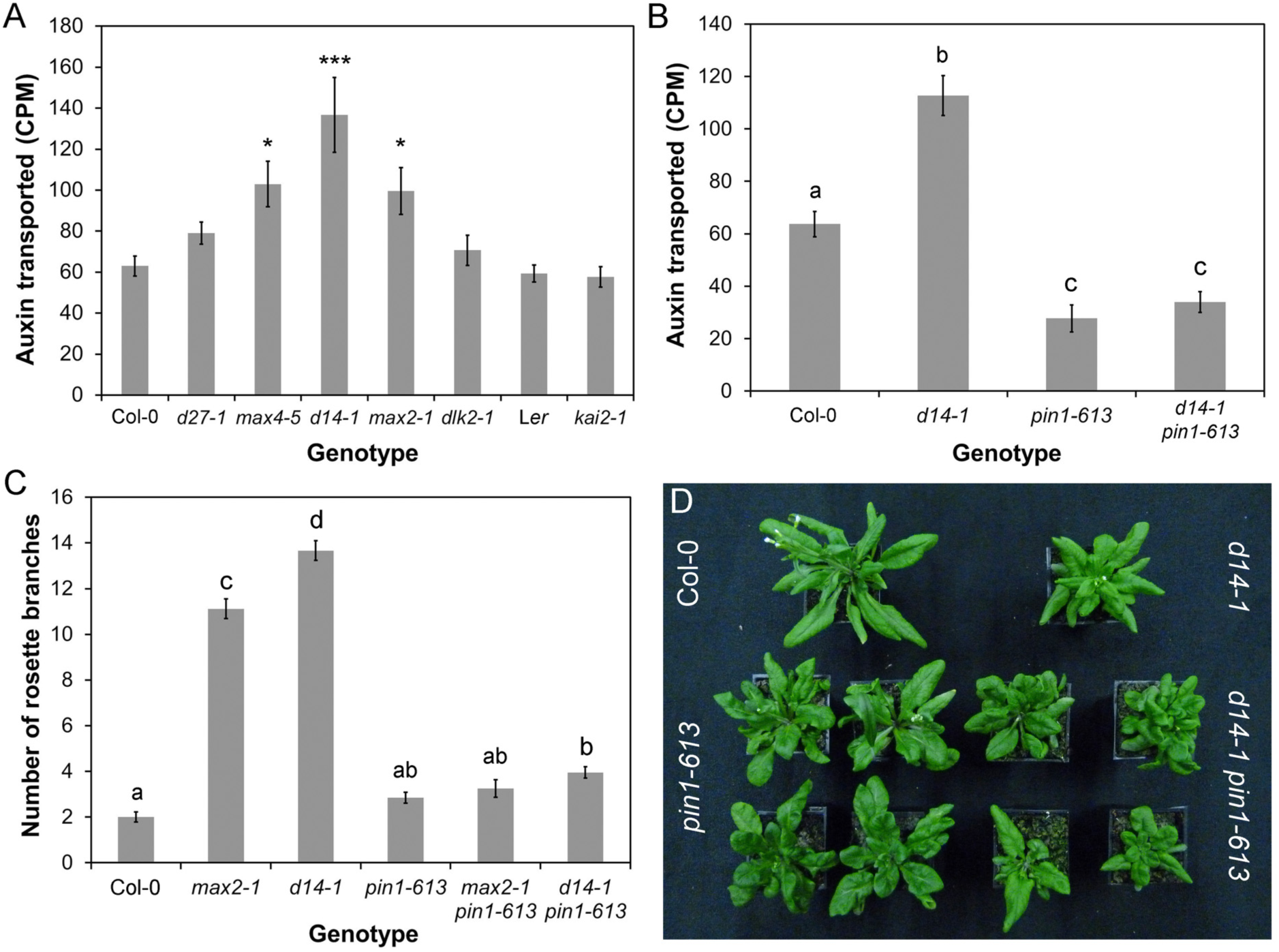
Canonical SL signalling affects stem auxin transport. **A)** Bulk auxin transport levels in candidate SL signalling mutants. The amount of radiolabelled auxin (assessed as counts per minute, CPM) transported in 6 hours through basal inflorescence internodes was measured in the indicated genotypes 6 weeks after germination, n=18-20, bars indicate s.e.m. Asterisks indicate genotypes that are significantly different from Col-0 (ANOVA, Dunnett’s test, * p<0.05, ** p<0.01, *** p<0.001). **B)** Effect of *pin1-613* mutation on bulk auxin transport in wild-type and *d14-1* mutant backgrounds. The amount of radiolabelled auxin (assessed as counts per minute, CPM) transported in 6 hours through basal inflorescence internodes was measured in the indicated genotypes 6 weeks after germination, n=18-22, bars indicate s.e.m.. Bars with the same letter are not significantly different from each other (ANOVA, Tukey HSD test). **C)** Rosette branching in *d14-1 pin1-613* and *max2-1 pin1-613* double mutants. The number of 1^st^ order rosette branches was measured at the proliferative arrest point of Col-0, n=15-34, bars indicate s.e.m.. Bars with the same letter are not significantly different from each other (ANOVA, Tukey HSD test). **D)** Morphology of rosette leaves in Col-0, *d14-1*, *pin1-613* and *d14-1 pin1-613*. Although lack of PIN1 causes severe effects on leaf morphology, the overall shape of *pin1-613* and *d14-1 pin1-613* leaves is still characteristic of their SL signalling status.

**Figure 8:**
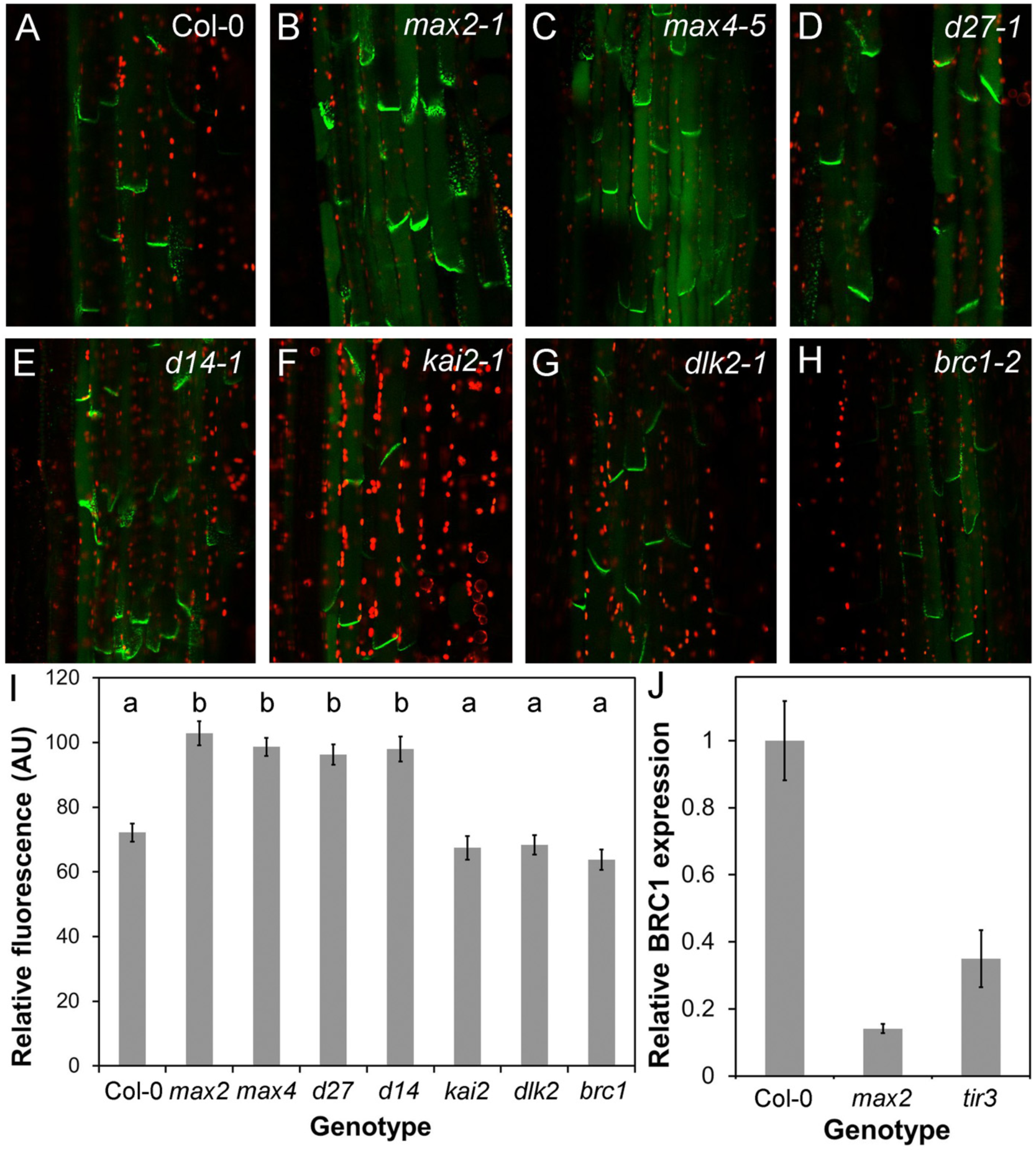
BRC1 and PIN1 act in parallel. **A-H)** PIN1:PIN1-GFP expression in wild-type, SL synthesis mutants and candidate SL signalling mutants. All images taken with identical settings, using hand sections through the basal inflorescence internode. **I)** Quantification of PIN1-GFP fluorescence on the basal plasma membrane in candidate SL signalling mutants, n=40 membranes per genotype (5 in each of 8 plants, except *max4-5* with 10 in each of 4 plants), bars indicate s.e.m. Bars with the same letter are not significantly different from each other (ANOVA, Tukey HSD,). **J)** Relative expression in *max2-1* and *tir3-101*,of *BRC1* in actively growing buds normalized to Col-0,, as assessed by qPCR. n=3 biological replicates per genotype, and 3 technical replicates per biological replicate. Error bars indicated s.e.m. of biological replicates.

Consistent with these observations, we have recently shown that increased SMXL7 levels are sufficient to increase auxin transport and PIN1 accumulation (Liang et al, 2016). We observed the same effect on auxin transport in *35:SMXL6*^Δ^*p-loop*^^-*YFP*, further demonstrating the equivalence in function of SMXL6 and SMXL7. Furthermore, we have also shown that loss of *smxl6*, *smxl7* and *smxl8* is causes a dramatic reduction in auxin transport and PIN1 levels in inflorescence stems (Figure S6A) (Soundappan et al, 2015). Thus, increased auxin transport and PIN1 levels in the inflorescence stem are consistent elements of the phenotypic syndrome caused by deficient SL signalling and resulting SMXL6 and SMXL7 accumulation.

Our previous results show that the increased shoot branching in *max* mutants is very likely caused at least in part by their altered PIN1 accumulation dynamics, such that increased steady state PIN1 levels and increased branching in the mutants both reflect a reduced rate of PIN1 removal from the plasma membrane (Shinohara et al, 2013; Prusinkiewicz et al, 2009; Bennett et al, 2006). The increased auxin transport seen in *d14-1* is suppressed in the *pin1-613* mutant background, consistent with the idea that it results from increased PIN1 accumulation (Figure 7B). The *d14-1 pin1-613*, *max1-1 pin1-613, max2-1 pin1-613* and *max3-9 pin1-613* also all have dramatically reduced shoot branching (Figure 7C) (Bennett et al, 2006). However, these data are difficult to interpret, since *pin1* mutants often fail to initiate axillary meristems, preventing an accurate assessment of axillary meristem activity (Wang Q. et al, 2014; Wang Y. et al, 2014).

With respect to leaf morphology, the *d14-1 pin1-613* and *max2-1 pin1-613* double mutants retain the characteristic leaf shapes of *d14-1* and *max2-1*, in addition to features characteristic of *pin1* such as leaf fusions (Figure 7D). This suggests that reduced PIN1 endocytosis is not the cause of the changes in leaf morphology caused by deficient SL signalling.

### BRC1 acts in parallel to PIN1

Our analysis suggests that BRC1 and PIN1 are plausible downstream targets of SL signaling, but in both cases, the evidence suggests they influence only a sub-set of SL-regulated phenotypes, in particular shoot branching. We therefore tested the relationship between BRC1 and PIN1 in the regulation of shoot branching. We assessed whether accumulation of PIN1 in the basal plasma membrane of stem xylem parenchyma cells was increased in *brc1-2*, but found that PIN1 levels are indistinguishable from wild-type (Figure 8A,H,I). Furthermore, we measured bulk auxin transport in *brc1-2 brc2-1*, and found that it is similar to wild-type, and significantly less than in *d14-1* (ANOVA, Tukey HSD, n=30, p<0.05) (Figure S6A). These data demonstrate that if BRC1 is involved in SL signalling, it does not act upstream of the regulation of PIN1 accumulation. We next tested whether *BRC1* expression is modulated by changes in PIN1 accumulation and/or auxin transport, i.e. whether *BRC1* is downstream of PIN1. The *max2* mutant has increased PIN1 accumulation and auxin transport, and reduced *BRC1* expression. Thus, we hypothesized that, if *BRC1* is downstream of PIN1, the *tir3* mutant, which has decreased PIN1 accumulation and auxin transport, ought to have increased levels of *BRC1* expression. However, we found that *BRC1* expression in *tir3* is strongly reduced, as in *max2* (Figure 8J). We thus conclude that BRC1 probably acts in parallel to PIN1 in the regulation of shoot branching.

## DISCUSSION

### SL perception in flowering plants

SLs are present, and can induce developmental effects, in charophyte algae and early diverging land plants. Whilst this implies the existence of SL signalling mechanism in these species, current evidence suggests that it must be markedly different from SL signalling in flowering plants. For instance, although present, MAX2 is apparently not involved in SL signalling in *Physcomitrella patens* (Challis et al, 2013; de Saint Germain et al, 2013a), and current phylogenetic analyses suggest that the SL receptor D14 appears to have evolved only within the vascular plants (Delaux et al, 2012; Waters et al, 2015). Conversely, KAI2-type proteins are found throughout land plants and charophyte algae, suggesting the existence of an ancient KAI2-mediated signalling pathway (which could be MAX2-independent) (Delaux et al, 2012; Bennett & Leyser, 2014). An interesting possibility therefore, is that SL signalling in early-diverging land plants is mediated by KAI2. Certainly, it appears possible that the vascular plant canonical SL signalling pathway has arisen by duplication and divergence of the ancestral KAI2 pathway, involving both the receptors (KAI2 and D14) and the immediate downstream targets (SMAX1 and SMXL7/D53), with MAX2 acting in both pathways (Bennett & Leyser, 2014). The possibility that KAI2 might be ancient SL receptor prompted us to examine whether KAI2 could be involved in SL responses in flowering plants. While it has previously been suggested that KAI2 acts mostly in seedlings and D14 later in shoot development (Waters et al, 2012a), we did find clear adult phenotypes for *kai2*. However, these were distinct from those found in *d14*, and all the phenotypes observed in the *max4* SL synthesis mutant are observed in *d14* alone. The *d14 kai2* double mutant resembled *max2*, showing that the additional adult phenotypes present in *max2* relative to *max4* most likely arise due to inactivity of the KAI2 signalling pathway in this mutant. KAI2 appears to have no role in SL signalling in the adult shoot in Arabidopsis, consistent with a significant body of work showing that KAI2 does not bind naturally occurring SLs and does not mediate apparent seedling responses to SLs (REFS). Where such responses have been attributed to KAI2, these are likely due to interaction with the non-natural enantiomers that are present in the widely used SL analog *rac*-GR24 (Scaffidi et al, 2013; Scaffidi et al, 2014). We also observed no strong phenotypes in the adult shoots of mutants in *DLK2*, the closest relative to *D14*, nor any reproducible enhancement of the *d14* or *kai2* phenotypes in double or triple mutants amongst these genes. Taken together these data suggest that D14 is the primary mediator of SL perception in the adult shoot in Arabidopsis.

### Direct targets of SL signalling

Recent reports have strongly implicated the chaperonin-like SMXL-family proteins as proteolytic targets of MAX2 in both KAI2- and D14-mediated signalling (Stanga et al, 2013; Zhou et al, 2013; Jiang et al, 2013; Soundappan et al, 2015; Wang et al, 2015). We show here that overexpression of a stabilized form of SMXL6 is sufficient to block SL responses in the adult shoot, further strengthening the idea that SMXL proteins are direct targets of SL signalling. Interestingly, our results suggest that some cross-activity between the KAI2 and D14 pathways is possible, because the stabilized form of SMXL6, like SMXL7 (Liang et al, 2016), is able to induce some *kai2*-like effects on leaf morphology when driven by the 35S promoter, in addition to the expected *d14*-like effects. This suggests either that high levels of SMXL6 can interfere with degradation of SMAX1, perhaps by titrating KAI2 or MAX2 out of the system, or that when ectopically expressed SMXL6 has some SMAX1-like activity.

Other direct targets of SL signalling have been proposed, and in this report, we have used comparative phenotypic analysis to assess their relative importance to SL responses. Morphological phenotypes can be influenced by many factors, making it difficult to determine whether similar phenotypes in different mutants have similar causes. To try to circumvent this we examined multiple adult shoot phenotypes using different genetic tools (including loss- and gain-of-function where possible) and used several different assays, including direct tests of SL sensitivity. Our results suggest that, contrary to previous suggestions, neither BES1 nor DELLA proteins fit the profile of an SL target in the regulation of shoot branching. DELLA proteins had only been implicated as SL targets on the basis of biochemical interaction with D14 (Nakamura et al, 2013), and previous reports in pea had suggested that they acted independently of SL in the regulation of internode elongation (de Saint Germain et al, 2013b). We did not find any evidence that DELLAs are SL targets in any aspect of development. BES1 was suggested as a SL target based on a mix of biochemical and phenotypic analysis, but using the highly pleiotropic *BES1-RNAi* line, and the original *bes1-d* line, which contains multiple segregating polymorphisms (Wang et al, 2013). Our analysis using back-crossed lines does not support any role for BES1 in shoot branching. Wang et al (2013) showed that in response to *rac*-GR24 treatment, BES1 can interact with MAX2, and is degraded in a MAX2-dependent manner. Given the apparent *rac*-GR24-insensitive hypocotyl elongation in *bes1-D,* it is possible that BES1 is a target of MAX2 in KL signalling. SL signalling and synthesis mutants do not have altered hypocotyl elongation, and in the hypocotyl, *rac*-GR24 primarily mimics the effects of KL signalling, and not SL signalling (Scaffidi et al, 2013; Scaffidi et al, 2014). More work is needed to test this possible role of BES1 in KL response.

In combination, our data suggest that with respect to the adult shoot phenotypes we assayed, the only direct targets of MAX2 are proteins of the SMXL6/7/8 clade. This is consistent with previous results showing that the *smxl6 smlx7 smxl8* triple mutant completely suppresses relevant aspects of the *max2* phenotype (Soundappan et al, 2015; Wang et al, 2015).

### Downstream targets of SL signalling

With regard to events further downstream, we have shown that BRC1/BRC2 and PIN1, but not SPL9/SPL15, are plausible SL signalling targets, but only in a sub-set of SL responses, primarily shoot branching.

The relationship between BRC1 and SL is complex. BRC1 has been widely described as acting downstream of SL based primarily on three observations. First, branching in *brc1* mutants and their equivalents in other species is SL resistant; second in double mutant combinations of SL and *brc1* mutants, branching levels are in some cases no higher than in the single mutants; and third *BRC1* expression levels are perturbed in SL mutant buds, and in pea *BRC1* transcription is up-regulated by SL in a cycloheximide-independent manner (Aguilar-Martinez et al, 2007; Braun et al, 2012; Minakuchi et al, 2010). However, while these data demonstrate the plausibility of *BRC1* acting as a downstream target of SL signalling, none is conclusive. SL insensitivity of *brc1* mutants is equally consistent with low BRC1 levels overcoming the effects of SL signalling via a parallel independent mechanism. Since most nodes produce an active branch in SL mutants, low additivity with other branching mutants is to be expected, and in any case is not universally observed. For example the *d14 brc1* double mutant can be more branchy than either parent (Chevalier et al, 2014). Similarly, the correlation between SL and *BRC1* transcription is not universal. For example, in rice, *FINE CULM1* (the *BRC1* paralogue) is not down-regulated in SL mutant buds and does not respond to SL treatment (Minakuchi et al, 2010; Arite et al, 2007). Furthermore, some of the effects of BRC1 on shoot branching might be the result of modulation of flowering time rather than direct effects on bud dormancy (Niwa et al, 2013; Tsuji et al, 2015). None of this precludes BRC1 being necessary for exogenous SL to inhibit shoot branching, but does mean the relationship cannot easily be explained as a simple linear one and more work is thus needed to clarify the exact role of BRC1 in branching control. For example, it is possible that *BRC1* transcription is up-regulated in dormant buds as a mechanism to stabilise their inactivity, rather than being required to impose dormancy *per se*.

Whether or not *BRC1* is a direct downstream target of SL signaling, it is clear that SL can affect shoot branching (and other shoot phenotypes) independently of BRC1. SL mutants can have stronger and different branching phenotypes than *brc* mutants (Figure 6) (Braun et al, 2013), and in maize SL deficiency increases branching even though the *BRC1* orthologue, *TB1*, is constitutively highly expressed (Guan et al, 2012). BRC1-independent SL activity could be mediated via effects on PIN1. There is good evidence that removal of PIN1 from the basal plasma membranes of xylem parenchyma cells is a direct primary response to SL addition (Shinohara et al, 2013). This mode of action has contributed to the development of the auxin transport canalization-based model for the regulation of shoot branching, and can explain the counter-intuitive observation that SLs can promote branching in auxin transport compromised genetic backgrounds (Shinohara et al, 2013). The PIN1 response has previously been shown to depend on MAX2, and here we show it is dependent on D14, but not KAI2 to or DLK2, as expected for a direct SL response. Consistent with this idea, we have previously shown that the over-accumulation of PIN1 in SL mutants can be completely suppressed in the *smxl6/7/8* triple mutant background (Soundappan et al, 2015), and that stabilization of SMXL7 is sufficient to increase PIN1 accumulation (Liang et al, 2016).. Interestingly, PIN1 accumulation is not affected in the *brc1 brc2* double mutant, demonstrating that altered PIN1 levels are not simply an indirect effect of increased branching, or a downstream effect of *BRC1/BRC2* down-regulation. Conversely, *BRC1* expression is not correlated with PIN1/auxin transport levels, suggesting that *BRC1* is not downstream of changes in PIN1, but rather acts in a parallel pathway.

### Strigolactone signaling and transcription

An interesting, and unresolved question, is whether SL signalling operates by modifying transcription of target genes, or is independent of transcription, or both, depending on the context and target. The current evidence for transcriptional regulation by SL signalling, even in the case of *BRC1*, is ambiguous. There are some changes in transcription upon treatment with *rac*-GR24, but the relevance of these is unclear (Mashiguchi et al, 2009). Conversely, we have previously shown the regulation of PIN1 by SL is independent of new translation (Shinohara et al, 2013). Proteins in the SMAX1 and SMXL6/7/8 clades have well-conserved EAR motifs, leading to an assumption that SMXL proteins modulate transcription through interactions with TOPLESS-family proteins (Zhou et al, 2013; Jiang et al, 2013; Smith & Li, 2014). Although SMXL7 can interact with TOPLESS-RELATED2 (TPR2) (Soundappan et al, 2015), the relevance of this interaction has not been established, and the EAR motif need not be involved in transcriptional regulation at all; there are other EAR-interacting proteins that could be partners for SMXL7 (Bennett & Leyser, 2014). Furthermore, we have recently demonstrated that SMXL7 lacking the EAR motif still possesses much, though not all of its functionality (Liang et al, 2016). This suggests that there could be separable EAR-dependent and -independent pathways downstream of SMXL7, which is consistent with our observation that neither altered PIN1 nor BRC1 levels can account for all the effects of SL in the adult shoot. For instance, the leaf shape phenotypes in *d14-1* are not suppressed by loss of *PIN1*, and loss of *BRC1/BRC2* does not cause any change in leaf morphology

One obvious possibility is that the other effects of SL might be mediated by changes in the localization and activity of other PIN family members, in different tissue contexts. Alternatively, these aspects of SL-signalling could be mediated by transcriptional or non-transcriptional downstream targets unrelated to those currently established for shoot branching. Thus, even though a core, canonical mechanism for SL signalling by D14/MAX2-mediated degradation of SMXL proteins is now well-defined, there remains much that we do not understand regarding the mechanism of SL action. Analysis of the broader effects of SL on plant development should yield valuable insights as to whether downstream effects are diverse, or whether there is a unified response mechanism.

## MATERIALS & METHODS

### Plant materials

The *max2-1* (Stirnberg et al, 2002), *max4-5*, *pin1-613* (Bennett et al, 2006), *tir3-101* (Prusinkiewicz et al, 2009) *d14-1*, *kai2-1*, *kai2-2*, *dlk2-1, dlk2-3* (Waters et al, 2012a), *d27-1* (Waters et al, 2012b), *brc1-2 brc2-1* (Aguilar-Martinez et al, 2007), *gai-t6 rga-t2 rgl1-1 rgl2-1 rgl3-1* (‘*della’*)(Feng et al, 2005), *gai-1* (Koornneef et al, 1985), *RGA:GFP-RGA* (Fu et al, 2003), *bes1-D* (Yin et al, 2002; González-García et al, 2011), *bes1-1* (He et al, 2005), *spl9-1 spl15-1* (Schwarz et al, 2008), *smxl6-4 smxl7-3 max2-1, smxl6-4 smxl7-3 smxl8-1 max2-1* (Soundappan et al, 2015) and *PIN1:PIN1-GFP* (Xu et al, 2006) lines have been described previously. *kai2-2*, *d14-1 kai2-2* and *d14-1 kai2-2 dlk2-3* each backcrossed 6 times into the Col-0 background were a kind gift from Mark Waters. Data are presented for *kai2-1* are in the Landsberg *erecta* background, except for Figure 8, where the *kai2-1* allele has been backcrossed into Col-0 background..Double mutants between lines were constructed using visible, fluorescent and selectable markers or by PCR genotyping as previously described (Waters et al, 2012a).

### Cloning

The *SMXL6* CDS was cloned into a pDONR221 entry vector (Life Technologies) (primers: ATGCCGACGCCGGTGACTACG and CCATATCACATCCACCTTCGCCG). The SMXL6^Δ^P-loop^^ variant, lacking amino acids 705-712 (FRGKTVVD), was made with the Q5 Site-Directed Mutagenesis Kit (NEB) (primers TACGTAACCGGTGAGTTATC and TTTGTCATCAAGGGAACAATG). SMXL6 and SMXL6^Δ^P-loop^^ entry clones were sub-cloned into a pEarlyGate101 destination vector, between the 35S promoter and a C-terminal YFP tag. The resultant constructs were transformed into the Col-0 or *max2-1* genetic background using the Agrobacterium floral dip method (Clough & Bent, 1998). Homozygous T3 lines were used for analyses.

### qPCR analysis

For *BRC1* gene expression analysis (Fig 8J), actively growing buds (>5mm) were harvested into liquid nitrogen. Total RNA was extracted using an RNeasy Plant Mini kit (Qiagen) and DNAse treated using the Turbo DNA-free kit (Ambion) as per manufacturer’s instructions, then quantified using a NanoDrop 1000. For cDNA synthesis, 500ng of total RNA was reverse transcribed with Superscript II (Thermo Fisher) according to manufacturer’s instructions. Quantification of transcript levels was carried out using SYBR Green reactions with 5ng cDNA in a 20µL volume on a Light Cycler 480 II (Roche) relative to the reference gene *UBQ10* (*UBIQUITIN 10*; At4g05320). Three technical replicates were run for each of three biological replicates. Expression levels were calculated using the Light Cycler 480 II software and the 2nd derivative maximum method assuming equal primer efficiencies. Primers: BRC1-F CTTAGTCAACTACAAACCGAACTCAT; BRC1-R GATCCGTAAACTGATGCTGCT; UBQ10-F CCACTTGGTCTTGCGTCTGC; UBQ10-R TCCGGTGAGAGTCTTCACGA.

### Plant growth conditions

Mature plants for analysis were grown on Levington’s F2 compost, under a standard 16 hr/8 hr light/dark cycle (22°C/18°C) in controlled environment rooms with light provided by white fluorescent tubes, (intensity ∼150 µMm^−2^s^−1^). For axenic growth, seeds were sterilised, and stratified at 4°C for several days. Seedlings were grown using ATS media (Wilson et al, 1990) with 1% sucrose, solidified with 0.8% plant agar, in 10cm square plates.

### Phenotypic measurements

The 7th leaf of each plant was marked with indelible marker at approximately 4 weeks post germination. These leaves were provisionally measured at 35 days post germination (dpg), and then measured again at 37 dpg to confirm that growth of these leaves was arrested. The maximum length and width of the leaf blade were measured, in addition to the length of the petiole (the petiole was not included in the blade length). Leaf senescence assays were performed as described by Stanga et al, (2013). Stem diameter, plant height, branch angles and branching levels were all measured at global proliferative arrest (approximately 7 weeks post germination), except where stated. Stem diameter was measured using digital calipers at the top and bottom of the basal inflorescence internode to obtain an average diameter. Height was measured using a ruler. Branch angle was measured by photographing the junction between the stem and the two basal-most cauline branches for each plant (or one, if there was only cauline node present). Using these images the angles between branch and stem using ImageJ was quantified for each plant, then averaged to obtain a single figure per plant. Standard branching level measurements were quantified as the number of 1^st^ order cauline and rosette inflorescences present on the plant. We also used a more sensitive decapitation-based assay to assess branching, in which plants are grown in short days to prolong the vegetative phase, generating more leaves and thus more axillary meristems (Greb et al, 2003). The plants are then shifted to long days to promote flowering and after the primary floral shoot reaches ∼10cm it is removed, activating inhibited axillary buds in the rosette. The number of elongated branches >1cm were counted 10 days after decapitation.

### Hormone response assays

Seeds were sterilized and stratified at 4°C for several days. The seeds were sown into 500ml jars (Weck, Germany) containing 60 ml ATS with 1% sucrose, solidified with 0.8% agar. For intact plant assays, plants were grown on media containing 5µM GR24 or an equivalent volume of acetone (solvent control) for 6 weeks, and branching was then measured. For excised nodal assays, plants were grown on plain ATS agar for ∼3 weeks, until bolting. Young nodes with buds <1.5mm in length were excised and placed between two agar blocks, to which hormones could be added independently (Chatfield et al, 2000). The growth of buds was then monitored daily over the following 10 days.

### Microscopy

For PIN1-GFP, GFP-RGA and SMXL6-YFP imaging in the shoot, hand sections were made through the vascular bundles of basal internodal stem segments of 6 week old plants, and the slices were then embedded in agar plates. For GFP-RGA GR24 treatments, stems were covered in ATS solution containing 5µM *rac*-GR24 or an equivalent volume of solvent control for 45 minutes before imaging. Images were taken using laser-scanning confocal microscopy using a Zeiss LSM700 imaging system with 20× water immersion lenses. Excitation was performed using 488 nm (15% laser power) and 555 nm (6%) lasers. Chloroplast autofluorescence was detected above 600 nm, and GFP/YFP fluorescence below 555 nm. The same settings for GFP/YFP detection were used within experiments for each line, except where stated. GFP quantification was performed on non-saturated images, using Zeiss ‘ZEN’ software. For PIN1-GFP, fluorescence intensity in the GFP channel was measured in four or five basal plasma membranes per sample, in at least 8 independent samples, except where stated. For RGA-GFP, fluorescence intensity in the GFP channel was measured in five nuclei per sample, in 8 independent samples per treatment.

For GFP-RGA imaging in the root, 7 day old seedlings were mounted on glass slides with 5µM *rac*-GR24 or an equivalent volume of solvent control in the mounting solution, then imaged after 45 minutes using a Zeiss LSM700 imaging system with a 20× lens. Excitation was performed using a 488 nm laser, and GFP fluorescence was detected below 555 nm. GFP quantification was performed on non-saturated images, using Zeiss ‘ZEN’ software. Fluorescence intensity in the GFP channel was measured in five nuclei per sample (2 in the epidermis and 1 each in the cortex, stele and root cap), in 12 independent samples per treatment.

For SMXL6-YFP imaging in the root, 3-5 day old seedlings were mounted on glass slides with 5µM *rac*-GR24, 5µM KAR1 or an equivalent volume of solvent control in the mounting solution, then imaged after 20 minutes using a Zeiss LSM780 imaging system with 20× lenses. For MG132 treatments, seedlings were pre-treated for 1 hour with 50µM MG132, then mounted as above. Excitation was performed using a 514 nm laser. YFP fluorescence was detected below 555 nm. The same settings for YFP detection were used within experiments for each line.

## ACKNOWLEDGEMENTS

This work was funded by the European Research Council (N° 294514 – EnCoDe), the Gatsby Foundation (GAT3272C), the UK Biotechnology and Biological Sciences Research Council via the European Research Area Plant Genomics programme (R1039101) and by the Chinese Government Scholarship PhD Program (Sichuan Agriculture University) to Y.L.

---

**Figure S1:**
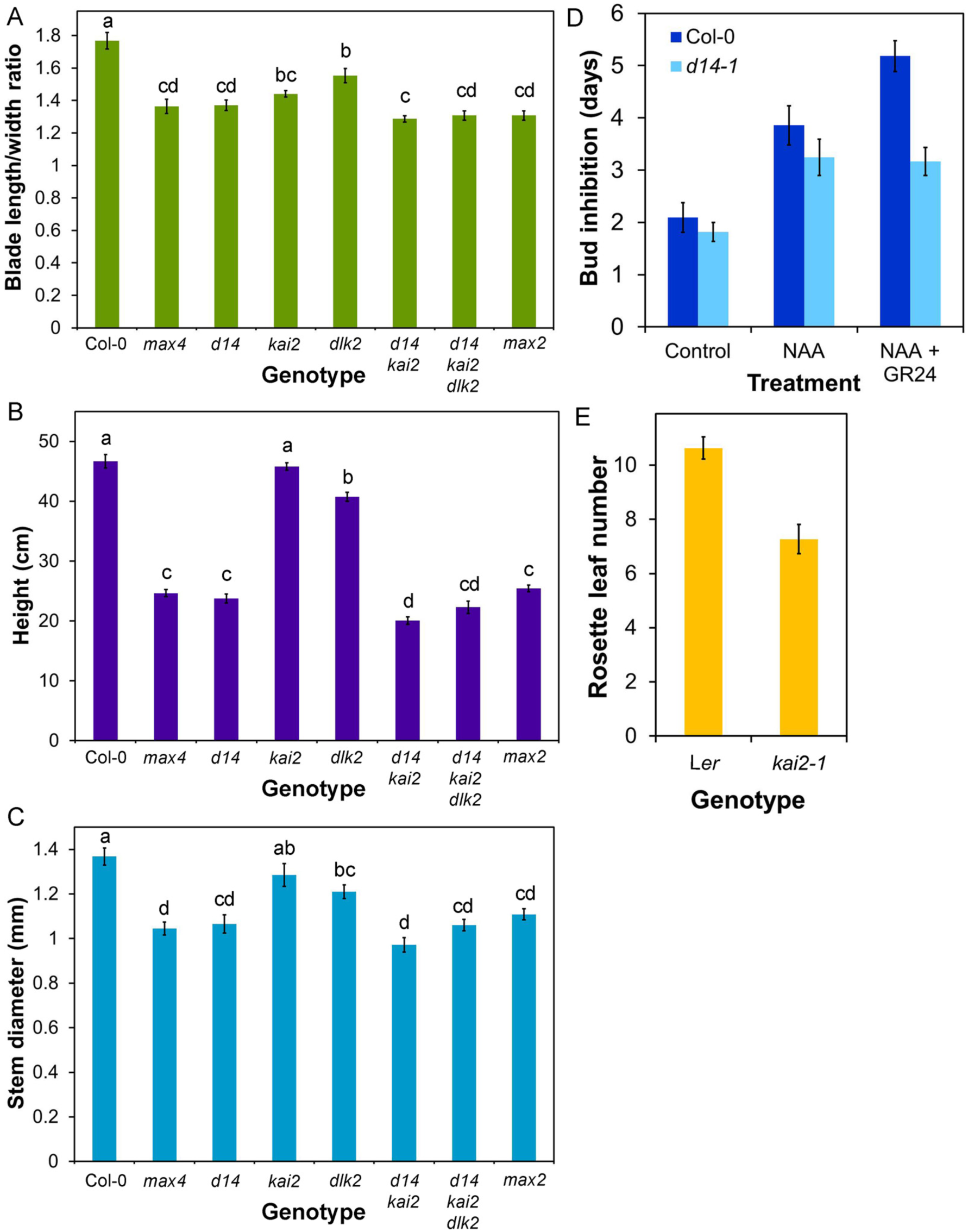
D14 mediates shoot SL signalling. **A)** Blade length:width ratios for candidate SL signalling mutants, calculated from the data in Figure 1D, n=10-12, bars indicate s.e.m. Bars with the same letters are not significantly different from each other (ANOVA + Tukey HSD test). **B)** Height (in cm) in candidate SL signalling mutants, n=10-12, bars indicate s.e.m. Bars with the same letters are not significantly different from each other (ANOVA + Tukey HSD test). **C)** Stem diameter (in mm) of the basal inflorescence internode in candidate SL signalling mutants, n=10-12, bars indicate s.e.m. Bars with the same letters are not significantly different from each other (ANOVA + Tukey HSD test). **D)** Growth responses of Col-0 and *d14-1* buds on excised nodal sections. Nodes were treated with either solvent control, 0.3µM NAA applied apically, or 0.3µM NAA apically + 5µM *rac*-GR24 basally. The average number of days that buds took to reach a length greater than 2mm is shown for each genotype and treatment, n=12-13 nodes per treatment, bars indicate s.e.m. **E)** Flowering time (measured as the number of rosette leaves produced before bolting) in *kai2-1* relative to L*er*, n=10-12, bars indicate s.e.m.

**Figure S2:**
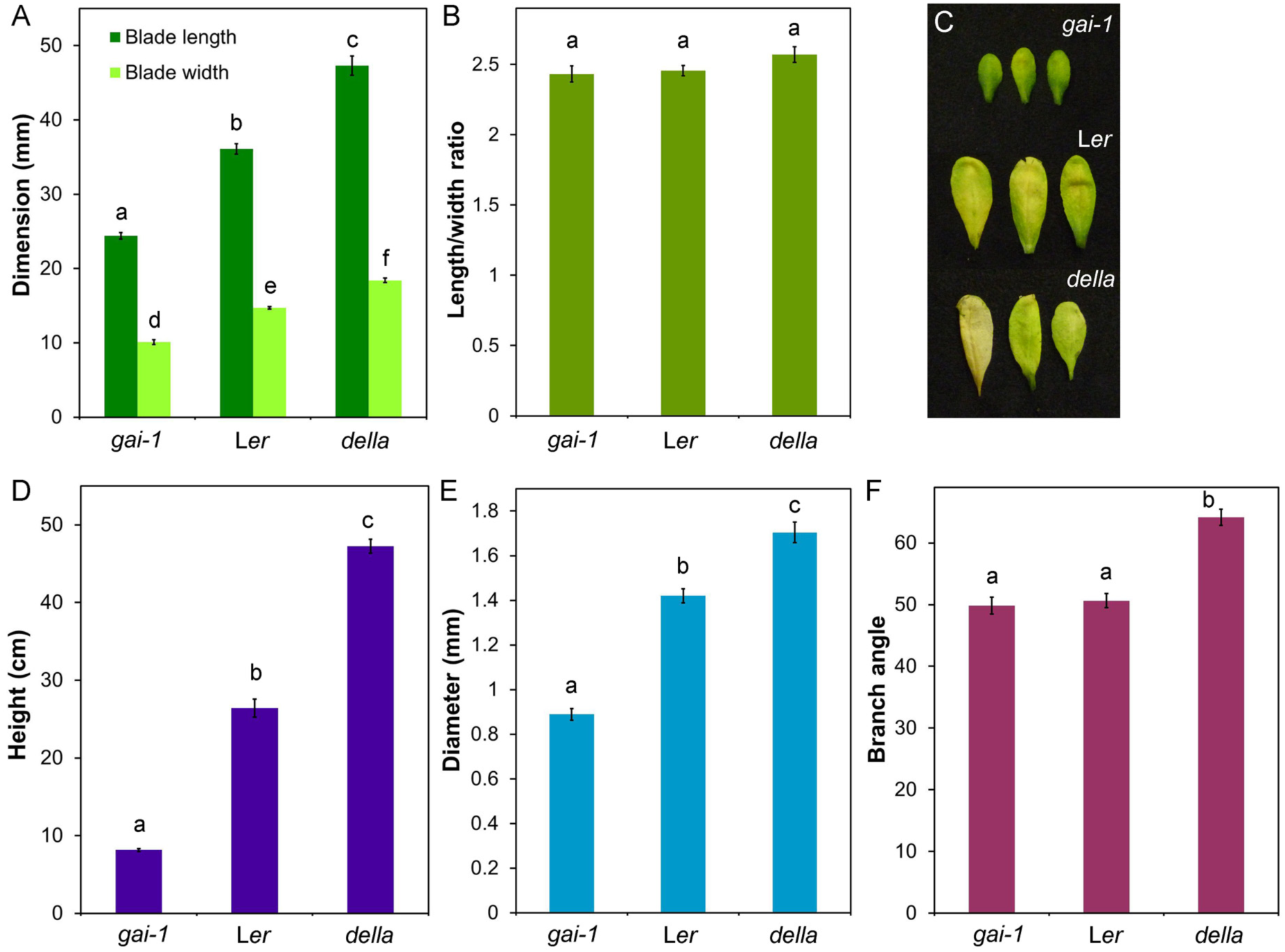
Role of KAI2 and DLK2 in shoot development. **A)** Leaf dimensions in Ler, *gai-1* and *gai-t6 rga-t2 rgl1-1 rgl2-1 rgl3-1* (*della*). Measurements were made on the 7^th^ rosette leaf, 35 days after germination. n=9-10, bars indicate s.e.m. Bars with the same letter are not significantly different from each other (ANOVA + Tukey HSD test). **B)** Leaf length:width ratio (including petiole) in L*er*, *gai-1* and *della.* n=9-10, bars indicate s.e.m. Bars with the same letter are not significantly different from each other (ANOVA + Tukey HSD test). **C)** Dark-induced senescence in L*er*, *gai-1* and *della* leaves from 5 week old plants. Leaves were wrapped in foil and imaged after 8 days. **D)** Plant stature in L*er*, *gai-1* and *della*, as assessed by the height of the main inflorescence stem (in cm), n=9-10, bars indicate s.e.m. Bars with the same letter are not significantly different from each other (ANOVA + Tukey HSD test). **E)** Stem diameter (in mm) of the basal inflorescence internode in L*er*, *gai-1* and *della*, as assessed by the height of the main inflorescence stem (in cm), n=9-10, bars indicate s.e.m. Bars with the same letter are not significantly different from each other (ANOVA + Tukey HSD test). **F)** Branch angle (in degrees) in L*er*, *gai-1* and *della*, n=9-10, bars indicate s.e.m. Bars with the same letter are not significantly different from each other (ANOVA + Tukey HSD test).

**Figure S3:**
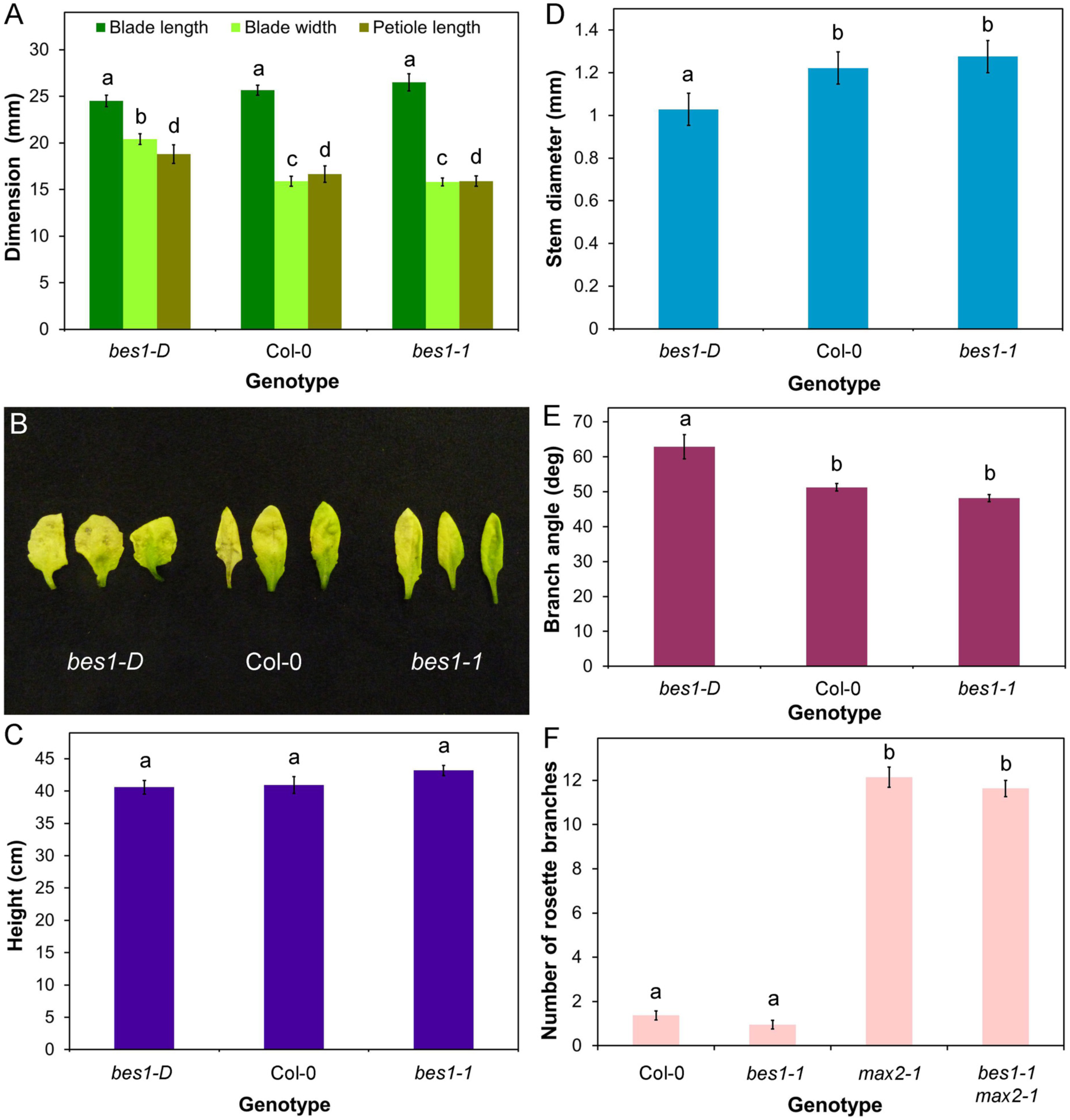
BES1 and SLs have different effects on shoot phenotype. **A)** Leaf dimensions in Col-0, *bes1-1* and *bes1-d*. Measurements were made on the 7^th^ rosette leaf, 35 days after germination. n=9-10, bars indicate s.e.m. Bars with the same letter are not significantly different from each other (ANOVA + Tukey HSD test). **B)** Dark-induced senescence in Col-0, *bes1-1* and *bes1-d* leaves from 5 week old plants. Leaves were wrapped in foil and imaged after 8 days. **C)** Plant stature in Col-0, *bes1-1* and *bes1-d*, as assessed by the height of the main inflorescence stem (in cm), n=9-10, bars indicate s.e.m. Bars with the same letter are not significantly different from each other (ANOVA + Tukey HSD test). **D)** Stem diameter (in mm) of the basal inflorescence internode in Col-0, *bes1-1* and *bes1-d*, as assessed by the height of the main inflorescence stem (in cm), n=9-10, bars indicate s.e.m. Bars with the same letter are not significantly different from each other (ANOVA + Tukey HSD test). **E)** Branch angle (in degrees) in Col-0, *bes1-1* and *bes1-d*, n=9-10, bars indicate s.e.m. Bars with the same letter are not significantly different from each other (ANOVA + Tukey HSD test). **F)** Numbers of primary rosette branches in long-day grown Col-0*, bes1-1*, *max2-1* and *bes1-1 max2-1.* Branching was measured at proliferative arrest, n=19-20, bars indicate s.e.m. Bars with the same letter are not significantly different from each other (ANOVA + Tukey HSD test).

**Figure S4:**
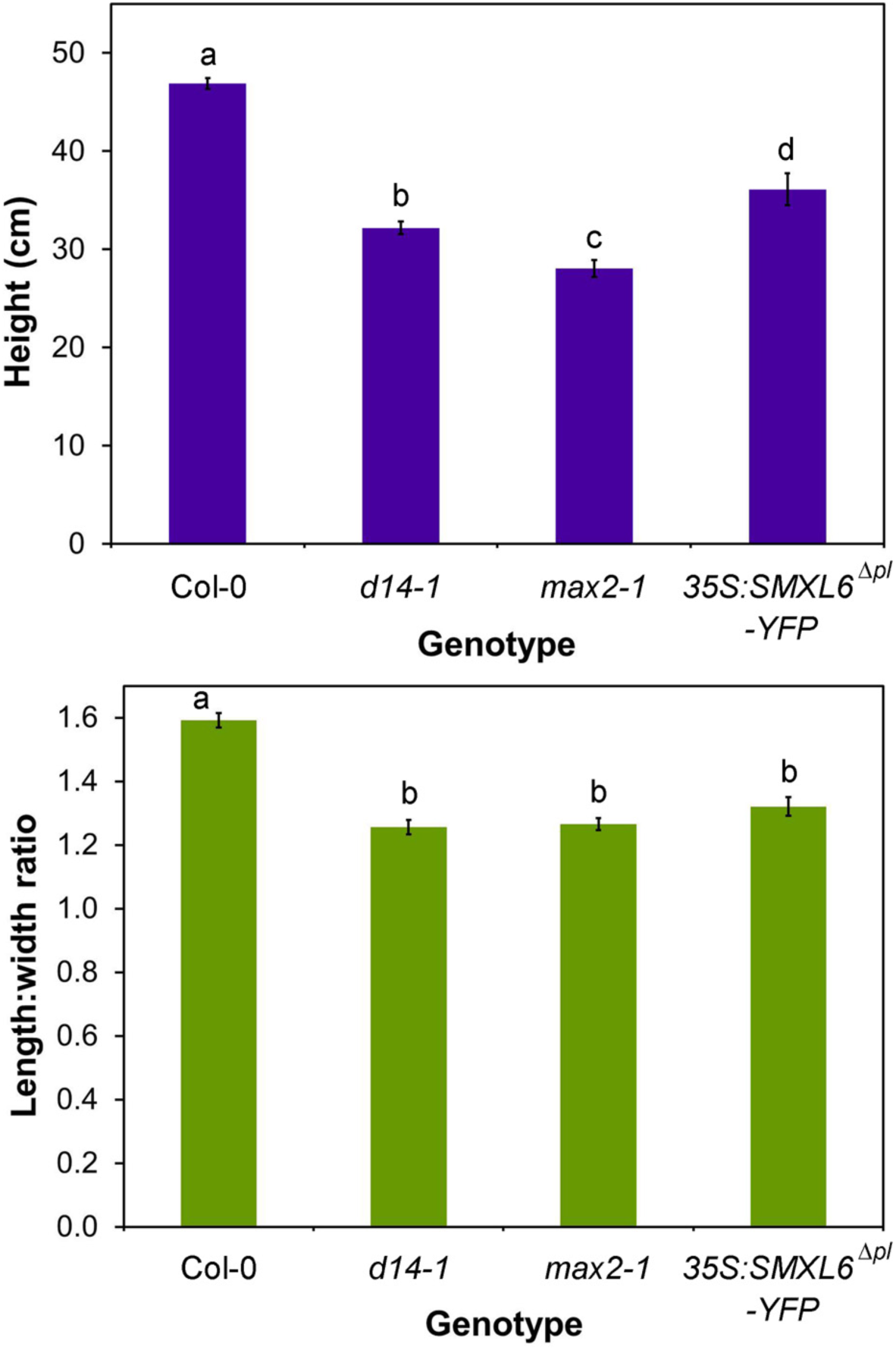
SMXL6 is functionally similar to SMXL7. **A)** Plant stature in Col-0*, d14-1*, *max2-1* and *35S:SMXL76*^Δ^*pl*^^-*YFP*, as assessed by the height of the main inflorescence stem (in cm), n=11-12, bars indicate s.e.m. Bars with the same letter are not significantly different from each other (ANOVA + Tukey HSD test). **B)** Leaf blade length:width ratio in Col-0*, d14-1*, *max2-1* and *35S:SMXL76*^Δ^*pl*^^-*YFP.* n=11-12, bars indicate s.e.m. Bars with the same letter are not significantly different from each other (ANOVA + Tukey HSD test).

**Figure S5:**
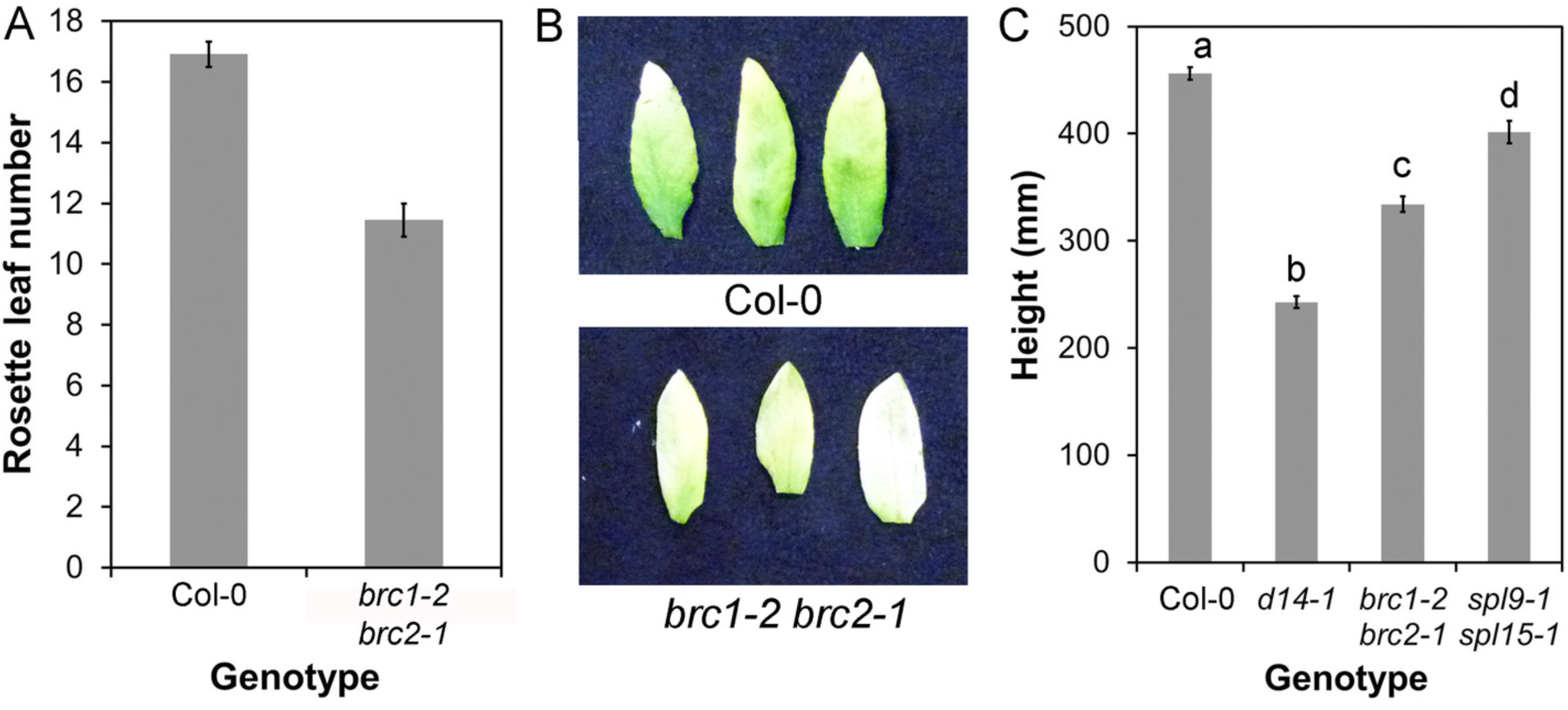
Phenotypic analysis of *brc1 brc2*. **A)** Flowering time in Col-0 and *brc1-2 brc2-1*, as assessed by rosette leaf number, n=11-12, bars indicate s.e.m. **B)** Dark-induced senescence phenotypes in Col-0 and *brc1-2 brc2-1*. Rosette leaves were wrapped in foil for 6 days then imaged. **C)** Final plant height in Col-0, *d14-1*, *brc1-2 brc2-1* and *spl9-1 spl15-1*. Height of the primary inflorescence was measured at proliferative arrest, n=11-12, bars indicate s.e.m.. Bars with different letters are significantly different from each other (ANOVA + Tukey HSD test).

**Figure S6:**
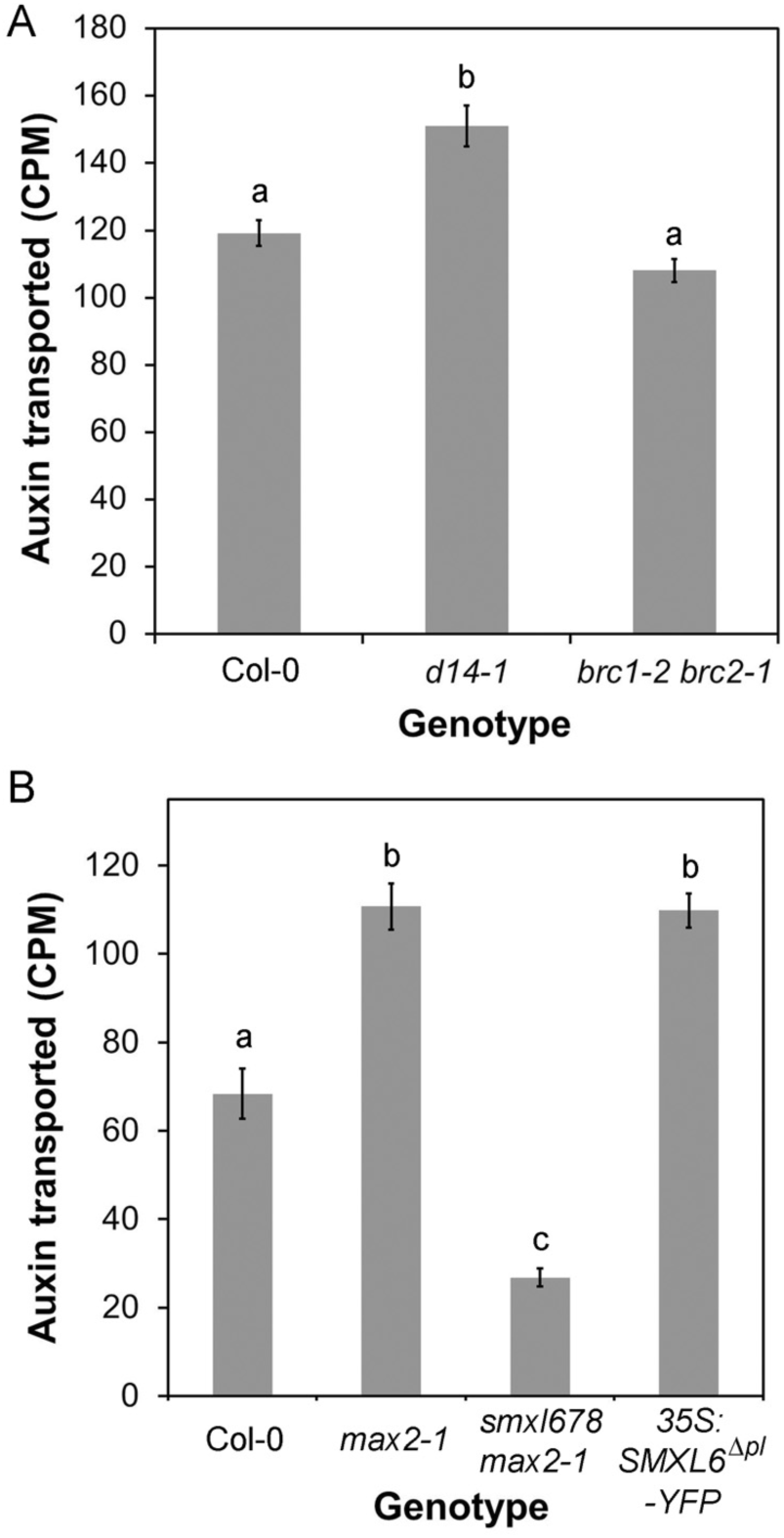
SL signalling and auxin transport. **A)** Bulk auxin transport through stem segments of in Col-0, *d14-1* and *brc1-2 brc2-1*. The amount of radiolabelled auxin (assessed as counts per minute, CPM) transported in 6 hours through basal inflorescence internodes was measured 6 weeks after germination, n=30, bars indicate s.e.m. Bars with the same letter are not significantly different from each other (ANOVA, Tukey HSD test). **B)** Bulk auxin transport through stem segments of Col-0, *max2-1, smxl6-4 smxl7-3 smxl8-1 max2-1* and *35S:SMXL6*^Δ^*p-loop*^^-*YFP*. The amount of radiolabelled auxin (assessed as counts per minute, CPM) transported in 6 hours through basal inflorescence internodes was measured 6 weeks after germination, n=30, bars indicate s.e.m. Bars with the same letter are not significantly different from each other (ANOVA, Tukey HSD test).

